# Nuclear localization and transactivation of SYS-1/β-catenin is the result of serial gene duplications and subfunctionalizations

**DOI:** 10.1101/2021.12.26.473689

**Authors:** Arielle K. Wolf, Lori C. Adams-Phillips, Amanda N. D. Adams, Albert J. Erives, Bryan T. Phillips

## Abstract

β-catenin is a multifunctional protein capable of mediating cell adhesion via E-cadherin and transactivation of target genes of the canonical Wnt signaling pathway. The nematode, *C. elegans* contains four paralogs of β-catenin which are highly specific in their functions. Though similar in overall structure, the four beta-catenins are functionally distinct, each regulating different aspects of development. Of the four, SYS-1 is a key player in Wnt dependent asymmetric cell division (ACD). In ACD, a polarized mother will give rise to a daughter with high nuclear SYS-1 and another with low nuclear SYS-1. Despite sequence dissimilarity, SYS-1 shares a close structural resemblance with human β-catenin where it retains an unstructured amino-terminus (NTD) and 12 armadillo repeats. Using existing genome sequence data from several nematode species, we find that the four β-catenin paralogs result from 3 sequential gene duplications and neofunctionalizations during nematode evolution. SYS-1, however, lacks an unstructured carboxyl-terminus (CTD) that is essential for human β-catenin transactivation processes. This work supports the hypothesis that SYS-1 compensated for the lack of CTD by acquiring novel transactivation domains with cryptic nuclear localization signals in the NTD and the first four armadillo repeats, as shown by transactivation assays in worms and yeast. Furthermore, SYS-1 regulatory domains are not localized to the NTD as in canonical β-catenin and instead spans the entire length of the protein. Truncating SYS-1 abolishes the classical SYS-1 nuclear asymmetry, resulting in daughter cells with symmetrical SYS-1 truncation localization. A screen for SYS-1 physical interactors followed by in vivo cell fate and SYS-1 localization analyses suggest that proper SYS-1 nuclear export is facilitated by XPO-1, while an interaction with IMB-3, an importin β-like protein, suggests import mechanisms. Interestingly, XPO-1 is especially required for lowering SYS-1 in the Wnt-unsignaled nucleus, suggesting a distinct mechanism for regulating asymmetric nuclear SYS-1. In summary, we provide insights on the mechanism of β-catenin evolution within nematodes and inform SYS-1 transactivation and nuclear transport.

## Introduction

Gene duplication events play an important role in the evolution of organisms by providing the raw genetic material for neofunctionalization (Katju and Bergthorsson, 2013). With the routine availability of whole genome sequence assemblies, it is now possible to systematically study the role of gene duplications in organismal evolution. The consequences of irreversible gene duplications through sub/neofunctionalizations are especially widespread when they impact transcriptional regulators because the evolution of new function, due in part to newly gained flexibility in protein sequence and resulting alterations to recognized cognate DNA sequences, can lead to a new cadre of downstream transcriptional events thus amplifying and rendering irreversible the effects of the original gene duplication. Here we show how three sequential gene duplications of an ancestral transcriptional regulator became irreversible in nematodes when they led to the generation of 4 essential sub/neofunctionalized loci.

One well studied and exemplar transcriptional regulator is β-catenin, also known as armadillo in *Drosophila*. In most animals, β-catenin homologs have long been known to be dual-functioned; they assist in cell adhesion by binding with E-cadherin and α-cadherin and also as a transactivator of the canonical Wnt signal transduction pathway (Figure 1A; Nelson and Nusse 2004). Reception of a Wnt ligand via the Frizzled receptor results in β-catenin stabilization and nuclear translocation. Nuclear β-catenin binds members of the LEF/TCF family of HMG transcription factors and activates the expression of TCF target genes. Thus, in the context of Wnt signaling, β-catenin regulates a wide array of cell fate decisions during metazoan development and adult tissue homeostasis while, in the context of cell adhesion, β-catenin also maintains epithelial integrity of a tissue.

**Figure 1.**
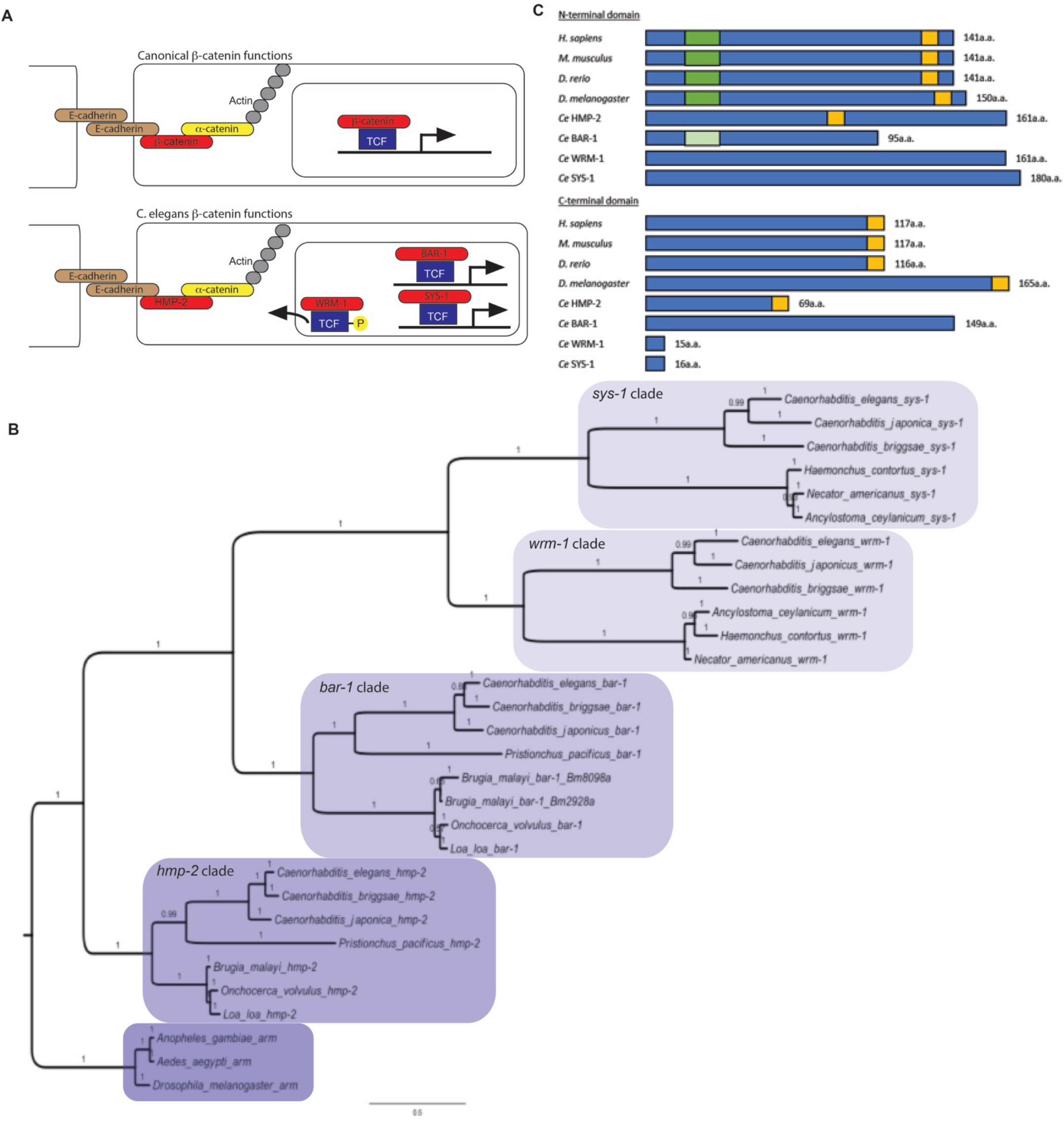
Serial neofunctionalization events led to SYS-1. **A.** Comparison between mammalian and *C. elegans* β-catenin function. Mammalian canonical β-catenin assists with adhesion (through the interactions with E-cadherin and α-cadherin) and transcriptional regulation (through the binding of TCF in the nucleus) (top). Nematode *C. elegans* contains four β-catenins with distinct and specialized roles in adhesion (HMP-2), TCF export (WRM-1) and transcriptional regulation via TCF/POP-1 (BAR-1 and SYS-1) (bottom). **B.** Bayesian MCMC support for a single family of nematode arm homologs that includes HMP-2, BAR-1, WRM-1, and SYS-1. Mixed amino acid models were sampled in the analysis, but the dataset supported only the Jones model. Posterior probabilities for branches are shown and 300,000 generations were found to be sufficient for convergence. **C.** Sequence conservation of β-catenin’s NTD and CTD. Conserved sites between *H. sapiens, M. musculus, D. rerio, D. melanogaster* β-catenins and *C. elegans* HMP-2 are in yellow. Conserved sites between *H. sapiens, M. musculus, D. rerio, D. melanogaster* and *C. elegans* BAR-1 are in green. Lighter shade indicates less sequence conservation.

β-catenin protein sequences are generally well conserved in both structure and function across different metazoans; for instance hydra β-catenin shares overall 62% identity to human β-catenin (Robertson and Lin, 2012; Schneider et al., 2003) and the sequence similarity is even higher in the central armadillo repeat region (Kidd et al., 2005). The amino terminal domain (NTD) as well as the carboxyl terminal domain (CTD) flanking the armadillo repeats are essential for cell adhesion and transcriptional regulation via Wnt signaling (Cox et al., 1999; Valenta et al., 2011). While the NTD facilitates the transcriptional role of β-catenin via recruitment of BCL-9 and subsequently Pygopus (Liu et al., 2008; Mosimann et al., 2009; Valenta et al., 2011) and the central armadillo repeats facilitate interaction with TCF (Stadeli et al., 2006), the CTD facilitates interactions between β-catenin and several chromatin-modifying factors including histones acetyltransferases such as CBP, the SWI/SNF factor BRG-1 and the mediator component MED12 (Barker et al., 2001; Hecht et al., 2000; Kim et al., 2006). Consistent with an essential role in transcriptional activation, the truncation of the CTD of *Drosophila* armadillo decreases its transcriptional activity compared to wild type (Cox et al., 1999). However, throughout evolution some β-catenin homologs have lost dual-functionality and became specialized in either cell-cell adhesion or transcriptional regulation (Chai et al., 2010; Korswagen et al., 2000). For example, the nematode *C. elegans* contains four homologs of β-catenin, HMP-2, BAR-1, WRM-1, and SYS-1 (Rocheleau et al., 1999; Kidd et al., 2005; Liu et al., 2008) and while the structure of the armadillo repeats is conserved amongst these β-catenin homologs, the N-terminal and C-terminal domains of *hmp-2, bar-1*, and *wrm-1* are highly divergent (Schneider et al., 2003) and the extent of SYS-1 sequence divergence from other β-catenins required its identification via functional analyses, rather than sequence similarity (Kidd et al., 2005, Miskowski et al,. 2001). This may have impacted the divergent functions represented by these β-catenins resulting in a separation of cellular functions between homologs.

Functional studies suggest that two *C. elegans* β-catenins, *hmp-2* and *bar-1*, are paralogs that divided the ancestral β-catenin functions of adhesion and transcriptional regulation between them. Consistent with a role as an adhesive β-catenin, HMP-2 localizes to adherens junctions and binds to E-cadherin HMR-1 but not the sole TCF homolog, POP-1 (Korswagen, 2002; Korswagen et al., 2000; Natarajan et al., 2001)(Fig 1A). The *hmp-2* mutant phenotype resembles loss of adherens junctions rather than *pop-1* mutants, suggesting that *hmp-2* is not required for Wnt signaling (Costa et al., 1998; Thorpe et al., 1997). Conversely, *bar-1*/β-catenin is a potent activator of transcription in the canonical Wnt/β-catenin signaling pathway. BAR-1 protein binds to POP-1/TCF using the conserved POP-1 N terminal β-catenin binding domain and activates transcription of Wnt target genes (Eisenmann et al., 1998)(Fig 1A). Furthermore, BAR-1 does not interact with E-cadherin or α-catenin, consistent with a conserved role in TCF activation rather than cell adhesion (Korswagen et al., 2000).

The third β-catenin, *wrm-1*, identified via sequence analysis (Rocheleau et al., 1997), functions as part of a Wnt/β-catenin pathway (termed the WβA pathway) that polarizes asymmetrically dividing mother cells during *C. elegans* development (Jackson and Eisenmann, 2012; for reviews, see Mizumoto and Sawa, 2007; Phillips and Kimble, 2009; Robertson and Lin, 2012; Lam and Phillips, 2017). A critical part of this pathway is POP-1/TCF regulation by WRM-1/β-catenin by directing POP-1 nuclear export and thus decreasing the ability of POP-1 to repress Wnt target gene transcription (Liu et al., 2008; Lo et al., 2004; Rocheleau et al., 1999; Yang et al., 2011; Zacharias et al., 2015). Therefore, compared to the typical transactivation mechanism seen in vertebrate β-catenin and BAR-1, WRM-1 controls TCF function by a novel subcellular localization mechanism (Fig 1A).

The fourth *C. elegans* β-catenin identified was SYS-1. Like WRM-1, SYS-1 functions positively in the Wnt ACD pathway, but the two β-catenins do not share the same molecular role. Instead, SYS-1 binds POP-1/TCF via the β-catenin binding domain and activates transcription (Kidd et al., 2005). In the absence of Wnt ligand, POP-1 mediates transcriptional repression through the interaction with histone deactylase I/HDA-1 and Groucho/UNC-37 (Calvo et al., 2001). Wnt activation stabilizes SYS-1 thus allowing SYS-1 to migrate into the nucleus. Current models suggest that, similar to vertebrate systems, binding of SYS-1 to POP-1 in the nucleus displaces UNC-37 thereby converting POP-1/SYS-1 complex into activators (Daniels and Weis, 2005). SYS-1 protein localization and regulation by the Wnt pathway is also similar to mammalian β-catenin (Baldwin et al., 2016; Baldwin and Phillips, 2014; Fuentealba et al., 2008; Gerhardt et al., 2015; Mila et al., 2015; Phillips et al., 2007; Vora and Phillips, 2015; Lam and Phillips 2017; Vora et al., 2020)(Fig 1A). Thus SYS-1 appears to function as a canonical β-catenin.

While the functions and structures of β-catenin homologs BAR-1, WRM-1, and HMP-2 are well studied in *C. elegans*, how SYS-1 structure relates to its function remains relatively elusive. The solved SYS-1 crystal structure, like human β-catenin, reveals 12 armadillo repeats that wrap in a superhelical fashion around a positively charged groove (Liu et al., 2008). The crystallized SYS-1/POP-1 complex shows a conserved “charged button” motif within this groove that is required for formation in both SYS-1/POP-1 and human β-catenin/TCF complexes. However, despite these structural, functional and regulatory similarities, SYS-1 was discovered by genetic screens (Miskowski et al., 2001; Siegfried and Kimble, 2002; Siegfried et al., 2004). SYS-1 was missed by sequence searches because it has little (~11%) sequence identity to human β-catenin (Kidd et al., 2005). Considering this poor sequence conservation yet significant functional and structural conservation, SYS-1 could be considered a potential β-catenin analog that may have evolved from a gene of unrelated function, i.e. SYS-1 could be considered a prime example of convergent molecular evolution. On the other hand, the crystal structures of both SYS-1 and the SYS-1/POP-1 complex display a high degree of conservation around the TCF binding site compared to human β-catenin and TCF extending even to identical salt bridges that anchor both complexes (Graham et al., 2000; Liu et al., 2008). Such a strong conservation suggests a degree of common ancestry between SYS-1 and other β-catenins, an idea we test below.

Through phylogenetic molecular sequence analysis, we found that SYS-1 is result of multiple sequential duplication and functionalization events from its ancestral β-catenin. We further found a role for transactivation in the NTD of SYS-1 and that the NTD is required for proper asymmetry cell division and WβA signaling. Considering the high structural and transcriptional functional conservation yet lack of sequence similarity between homologs, we propose SYS-1 to be a product of convergent molecular evolution. We follow these phylogenetic studies with a structure/function approach to inform SYS-1 transactivation and nuclear localization that identify transactivation domains and cryptic nuclear localization sequences.

## Results

### Serial neofunctionalization events led to *sys-1: arm→bar-1→wrm-1→sys-1*

In order to examine the ancestral history of SYS-1, we applied a neighbor joining analysis to closely related β-catenin sequences along with outgroup sequences containing bifunctional *arm* orthologs from the dipteran genera *Drosophila, Anopheles*, and *Aedes*. We found that *hmp-2* and *bar-1 genes* are the result of a duplication of *Drosophila* β-*catenin/armadillo* (*arm*) with *bar-1* and *hmp-2* present across various nematode species (Figure 1B). When compared to the outgroup we concluded that the BAR-1 sequence undergoes greater change relative to *hmp-2* (Figure 1B; Supp. Figure 1,2). This signature is consistent with functionalization of the *bar-1* paralog and is consistent with adhesive functions of HMP-2 constraining the evolution to a greater extent than the transcriptional regulation activity of BAR-1. This loss of constraint may have enabled further duplications and functionalization of WRM-1 and SYS-1.

WRM-1 is involved in facilitating POP-1 export through a phosphorylation event with LIT-1. WRM-1 sequence shows a weak similarity with BAR-1 and from our phylogenetic analysis we found that *wrm-1* is a result of a duplication and further functionalized paralog of *bar-1* (Figure 1B; Supp. Figure 3). To understand the timing of this duplication and to rule out the possibility of horizontal gene transfer in the origin of *wrm-1*, we searched GenBank for the most closely related sequences. We find that the most closely related sequences after those from the *Caenorhabditis* genus (order RHABDITIDA) were nematode sequences from three genera in the sister order STRONGYLIDA (Supp. Figure 4). Sequence alignment and phylogenetic analysis showed that all related sequences were part of a *wrm-1* clade.

SYS-1 was originally identified from a forward genetic screen as required for the Wnt-signaled fate during ACD and subsequently identified as a POP-1 interactor. Due to the low sequence similarity to human β-catenin and BAR-1, we hypothesized that SYS-1 is was a result of a gene duplication and neofunctionalization event from *wrm-1*. Phylogenetic analysis was conducted on all *sys-1* sequences. Similar to *wrm-1, sys-1* was identified in the RHABDITIDA and STRONGYLIDA nematode orders (Supp. Figure 5), indicating its presence in their last common ancestor. Next, we took the *sys-1* sequences from the STRONGYLIDA clade and searched for nematode homologs in GenBank. In addition to the SYS-1 sequences the most closely related sequence was the *wrm-1* gene from *Haemonchus contortus* with an E-value just under e-10. This result is consistent with a closest homologous and detectable relationship of *sys-1* to the *wrm-1* paralog. A representative alignment between the predicted protein sequences from *sys-1* and *wrm-1* of *Haemonchus contortus*, shows that the while the E-value (8e-10) and percent identity (169/757 or 22%) are both modest, the percent similarity (313/757 or 41%) and percent coverage (88% of *sys-1*) are not inconsistent with homology. This sequence homology can be seen best in the alignment of the primary peptide sequences predicted by the *wrm-1* and *sys-1* loci of *Haemonchus contortus* (Supp. Figure 5,6).

These data suggest that the *sys-1* gene represents a divergent homolog caused by three serial neofunctionalizations of the metazoan *armadillo* gene during nematode evolution. In order to seek further support or contradiction of this view, we used a Bayesian Markov chain Monte Carlo package (MrBayes 3.2) to sample model space and infer a phylogenetic tree using Bayesian inference. All of the sequences were aligned using ClustalW and analyzed. We find support for a single united family of Armadillo proteins such that the most recent family of *sys-1* represents the lineage with the most neo-functionalized (fast-evolving) stems going back in time to the original duplication of the ecdysozoan *arm* gene into *hmp-2* and *bar-1* (Figure 1B).

Since the N-terminal and C-terminal domains of β-catenin have been shown to be essential for cell adhesion and transcriptional regulation, we next sought to determine the probable presence of such domains within the four *C. elegans* β-catenin homologs. Using the central Armadillo repeat-containing region as a guide, we aligned the N-terminal domain or the C-terminal domain respectively with known N-/C-terminal domains of β-catenin sequences from *H. sapiens, M. musculus, D. rerio*, and *D. melanogaster* and found conserved domains in the N- and C-terminal regions of HMP-2 which were absent in BAR-1, WRM-1, and SYS-1. BAR-1 contained a slightly conserved region that was absent in *hmp-2* (Figure 1C). While the absence of a CTD in WRM-1 is consistent with the loss of transactivation function of this β-catenin (Fig 1A), the absence of this domain was unexpected for SYS-1, which has been shown to transactivate target genes in nematodes, yeast and human cells

We conclude that *sys-1 is* the most divergent of the *C. elegans* β-catenin homologs, a result of multiple duplications and neofunctionalization events, and that *hmp-2* is closer in sequence similarity to the ancestral β-catenin compared to the other three homologs (Figure 1C, Supp. Figure 7). Given this evolutionary arrangement, these data indicate that the SYS-1 precursor had lost conserved domains in the C terminus that facilitate the transcriptional activation function in other β-catenins by the last common ancestor with WRM-1, suggesting this function is contained within residues located elsewhere in the SYS-1 sequence, an idea we test below.

### The SYS-1 NTD and R1-R4 are minimal transactivation domains in yeast one-hybrid assay

In order to identify the transactivation domain of SYS-1, we created various truncations of the SYS-1 protein and tested for transcriptional activity in yeast. Truncation sites were based on the known SYS-1/POP-1 interaction site (R5-8) as well as the predicted unstructured NTD domain and on the size of each individual armadillo repeat, all of which were gleaned from the SYS-1 and the SYS-1/POP-1 crystal structures (Liu et al., 2008). We tested the transactivation property of each truncation using the yeast one-hybrid (Y-1-H) assay (Ouwerkerk and Meijer, 2001) (Figure 2A). Among the truncations made (Supp. Figure 8), we found three truncations that exhibited varying degrees of transcriptional activation of the β-galactosidase target gene (Figure 2B). When compared to full length SYS-1, NTD-R4 exhibited the strongest transcriptional response followed by R1-R4, and the NTD. Other truncations exhibited no transcriptional activity as quantified by ONPG (Figure 2B) or X-gal assays above empty vector background (Figure 2B-C, Supp. Figure 9). Taking advantage of the presence of an additional LexA operator upstream of HIS3 gene, a second assay was performed on histidine deficient media. Consistent with the results obtained from the X-gal plates, only full-length SYS-1, NTD-R4, R1-R4, and the NTD grew on -HIS+3AT plates (Figure 2D).

**Figure 2.**
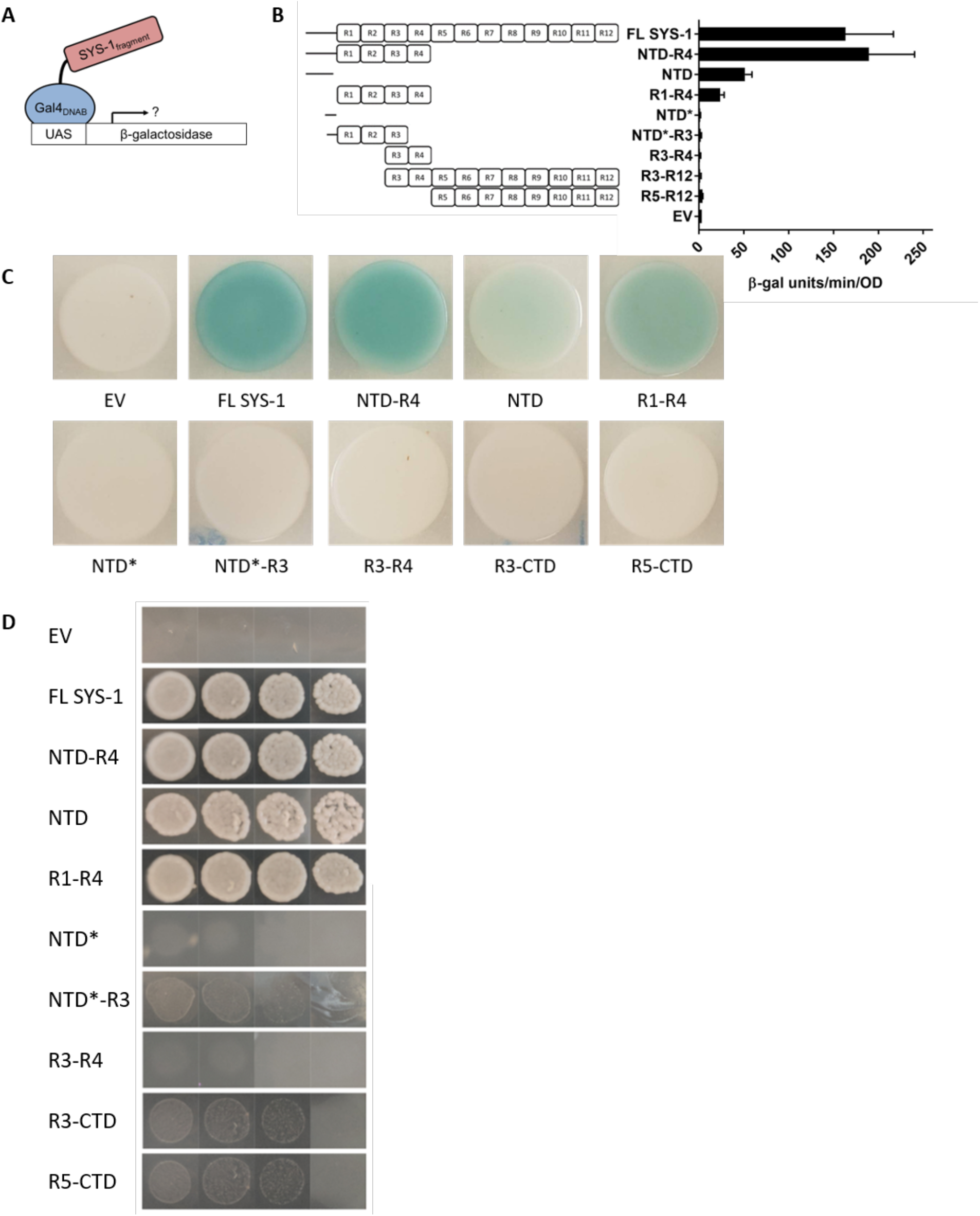
The SYS-1 NTD and R1-R4 are minimal transactivation domains in yeast. (A) Yeast one-hybrid used identify transactivators. Briefly, a LexA-SYS-1 truncations fusion protein were expressed allowing for effective binding to the LexA operator allowing testing for SYS-1 fragments transactivation activity. (B) ONPG assay on various truncated SYS-1 constructs. ONPG results from SYS-1 truncations were normalized by subtracting each reading from the empty vector readings. Data were shown as mean±SD. (C) X gal assay on Y-1-H constructs done on on plates containing X-gal (produces blue colonies upon transactivation activity) and - TRP media (selection media for plasmids containing SYS-1 truncations). (D) Serial-dilution of Y-1-H constructs used in ONPG assay. Yeast cultures were OD at 0.5 and undergoes 3 subsequent 1:10 serial dilutions before plating on respective plates. Growth assay were done on -HIS-TRP media to test for proper expression of SYS-1 truncations. 3AT were added to restrict growth due to the leaky HIS gene.

ortho-Nitrophenyl-β-galactoside (ONPG) assays (Reynolds et al., 2001; Mockli and Auerbach, 2004) were carried out to further verify and quantify transcriptional activity. From the ONPG assay, we found that both full-length SYS-1 (163 r.u.) and NTD-R4 (190 r.u.) are strong activators, followed by R1-R4 (51 r.u.), and NTD (24 r.u.). These collective results from Y-1-H demonstrates the role of NTD and R1-R4 as separable minimal transactivators in SYS-1 (Figure 2B). Though some coactivators bind the R1-2 armadillo repeats in other systems (Stadeli et al., 2006), these interactions do not extend into the unstructured NTD of β-catenin we identified here. Conversely, no transactivation activity was identified in the CTD, which is a transactivation domain in other systems (Takemaru and Moon, 2000). These data suggest that SYS-1 contains novel transactivation domain architecture that include two separable domains in the N-terminal half of SYS-1 (NTD and R1-R4) and that the conserved C-terminal transactivation activity found in vertebrate β-catenin has been lost in SYS-1.

### SYS-1 NTD and R1-R4 acts as a minimal transactivators in sensitized background in *C. elegans*

The somatic gonadal precursor (SGP) cells are the progenitor to the distal tip cells (DTCs) and the differentiation of the SGP to the DTC is governed by the asymmetric division of the SGPs (Kimble and Hirsh, 1979). Two SGPs, Z1 and Z4 each divide asymmetrically giving rise to a distal daughter with high nuclear SYS-1 and a proximal daughter with low nuclear SYS-1 (Phillips et al., 2007). Each distal daughter will give rise to a DTC, while the proximal daughters give rise to either anchor cells or ventral uterine cells (Figure 3A). Perturbations in the WβA pathway such as SYS-1 RNAi or overexpression of SYS-1 often causes the loss or increase, respectively, of DTCs within the nematode (Kidd et al., 2005).

**Figure 3.**
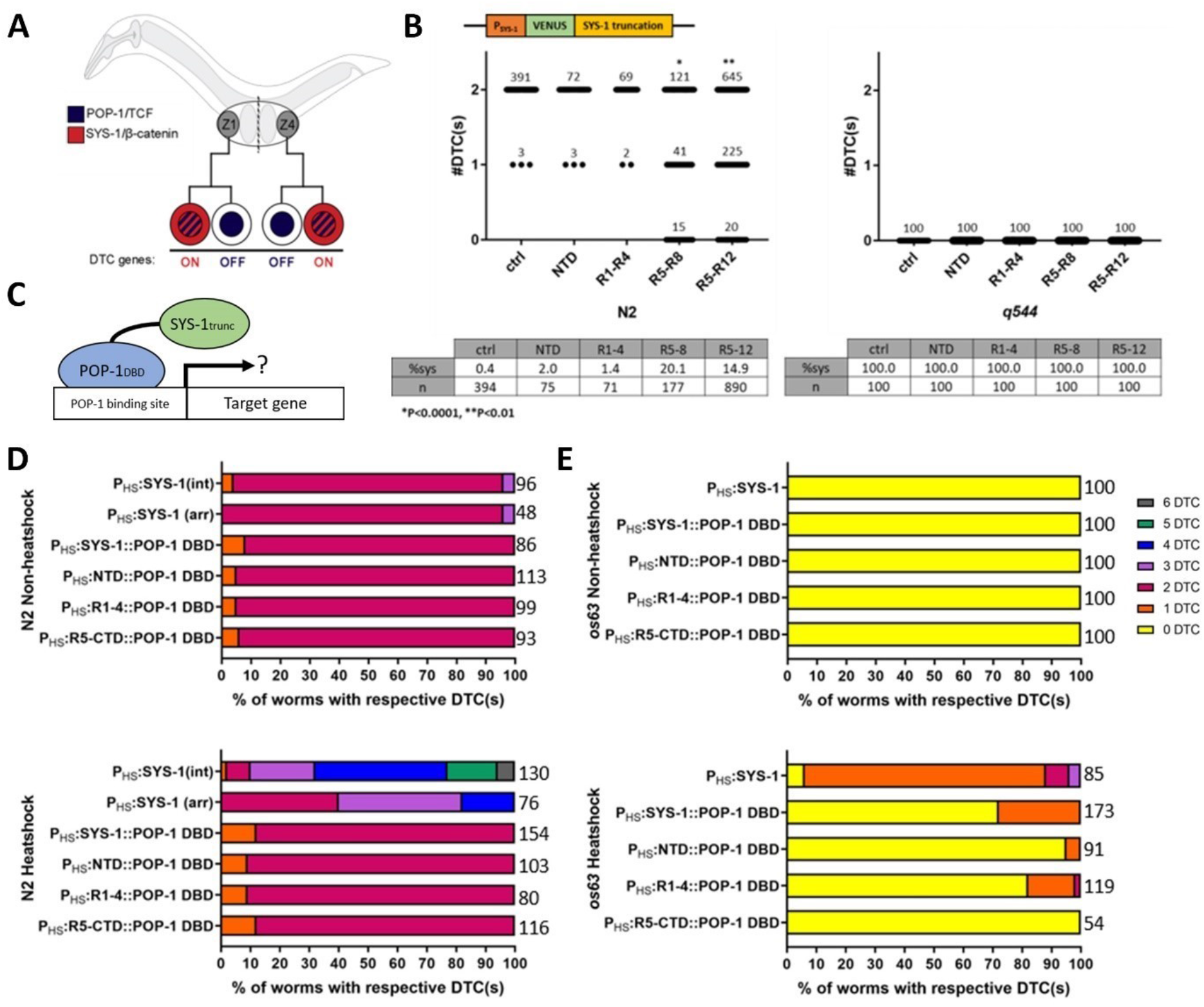
NTD and R1-R4 act as minimal transactivators in sensitized background in *C. elegans*. (A) WβA in DTC specification. (B) DTC counts on worms of SYS-1 truncations on extrachromosomal array in N2 (left) and q544 (right) backgrounds. *% Sys* was calculated based on the percentage of SGP failed to produce DTC. (C) Worm-1-hybrid for identifying SYS-1 truncations transactivation activity. POP-1 DBD fusion protein is used to tether SYS-1 truncations onto POP-1 DNA binding site for activation. DTC counts on worms of SYS-1 truncation constructs on extrachromosomal array in heatshock (right) or non-heatshock (left) N2 (D) and os63 (E) backgrounds.

To test if the expression of SYS-1 NTD or R1-R4 fragments into *C. elegans* would induce a cell fate change in the SGP lineage resulting in extra DTCs, we created transgenics containing Venus tagged SYS-1 truncations in wild type (N2) worms containing a DTC marker (P_lag-2_::GFP). Transgenics were synchronized and DTCs were counted at the early L3 larval stage. We observed no increase in DTC in the two truncations (NTD and R1-R4) that were shown as minimal transactivators from Y-1-H assay when compared to controls (Figure 3B left). We reasoned that the role of these truncations in regulating cell fate changes could be masked by the presence of endogenous full-length SYS-1, thus requiring a higher threshold for truncations to affect the changes of cell fate. To lower the threshold required for cell fate changes, we performed the same experiment in a *sys-1* hypomorph, *q544*. The *sys-1(q544)* strain, which produces no DTCs when maintained in 20°C, was used as a sensitized background for detecting NTD and R1-R4 transactivation. However, DTC counts from SYS-1 truncations also failed to suppress the *q544* mutant phenotype (Figure 3B right). Interestingly, DTC quantitation of strains containing the two non-transactivating truncations (R5-8 and R5-12) showed a statistically significant decrease of DTC counts in the N2 background. R5-R8 (~20% Sys, here defined as the percentage of SGPs that fail to give rise to a DTC) and R5-R12 (~15% Sys) showed a significant reduction of DTC when compared to controls (Figure 3B left). Since both the R5-8 and R5-12 fragments include the POP-1 interacting domain of SYS-1, it is likely that these SYS-1 fragments are binding and sequestering POP-1, leading to loss of POP-1 transactivation in the DTC lineage and subsequent DTC loss.

We thus hypothesized that, since both NTD and R1-4 fragments lack the POP-1 binding domain, the lack of symmetric divisions resulting in duplications of the DTC lineage observed in NTD or R1-R4 expressing animals were due to the inability of the truncations to functionally interact with POP-1, thereby preventing SYS-1 truncations from effectively recruiting coactivators to the regulatory region of Wnt target genes. To test this hypothesis, we created a system similar to Y-1-H in *C. elegans*, which we named “worm one-hybrid” (W-1-H). In W-1-H, the POP-1 DNA binding domain was fused to SYS-1 truncations to bring SYS-1 truncations in proximity with POP-1 target gene regulatory regions (Figure 3C). Conditional expression using the heatshock promoter was deemed necessary since we suspected that constitutive POP-1 target gene activation would be deleterious to worm development. DTC counts were performed on transgenics expressing truncations in N2 or sensitized *sys-1(os63)* background. *os63* hypomorphs were used instead of *q544* in these experiments as *q544* is suppressed by heatshock. While no increases in DTC number were seen in the N2 background (Figure 3D), when *os63* homozygotes were heatshocked, strains expressing full length SYS-1, NTD and R1-R4 fused to POP-1 DNA binding domain partially suppressed the *os63* phenotype; all showed an increase in DTC number in the sensitized *os63* background, with approximately 26% (FL SYS-1+POP-1 DBD), 5% (NTD+POP-1 DBD) and 18% (R1-4+POP-1 DBD) of individuals showing an increase in at least 1 DTC (Figure 3E). As expected, no effects on DTC specification were seen with worms expressing SYS-1 R5-CTD fused to the POP-1 DBD (Figure 3E). These data suggest that NTD and R1-R4 are indeed minimal activators when the truncations are brought to close proximity with target gene promoter sites.

The low percentage of presence of DTC rescue of the *os63* background after fusion expression may be caused by either mosaicism of the extrachromosomal arrays or the partial disruption of SYS-1 interactions of SYS-1 due to the addition of the POP-1 DNA binding domain. To test the role of mosaic transgene expression, we compared the DTC counts between integrated transgenic and extrachromosomal array transgenics of full-length SYS-1 under the heatshock promoter. We found that, in the N2 background, integrated full-length SYS-1 had more penetrant effects on additional DTC specification (90% of individuals exhibited more than two DTCs) compared to 60% transgenic worms expressing extrachromosomal array (~60%) (Figure 3D, Figure 3E). This suggests mosaicism may be a contributing factor in the low increase of DTCs of the worm one-hybrid experiments.

### SYS-1 truncations reveal SYS-1 nuclear localization domains

In addition to interactions with transactivating partners, β-catenin must also localize to the nucleus in order to activate target gene expression. However, the domains required for β-catenin nuclear localization are unknown and the β-catenin nuclear localization mechanism remain controversial (Anthony et al., 2020). We sought to identify the presence and location of possible NLS sites on SYS-1 through in vivo localization experiments by creating transgenics containing VENUS-tagged SYS-1 truncations protein expressed under the *sys-1* endogenous promoter and examined the resulting expression in the vulval precursor cells (VPCs). Like many anterior-posterior divisions in *C. elegans* development, the VPCs divide asymmetrically under the control of the WβA pathway. Current models suggest that a central Wnt source polarizes two central VPCs, each producing daughter cells with high nuclear SYS-1 in the proximal daughters and a low nuclear SYS-1 in the distal daughters (Phillips et al., 2007). Thus, VPC ACD results in two daughter pairs with mirror-image asymmetry expression in the vulva (Figure 4A), similar to the DTC lineage except the proximal VPC daughters represent the signaled cell fates instead of the distal SGP daughters.

**Figure 4.**
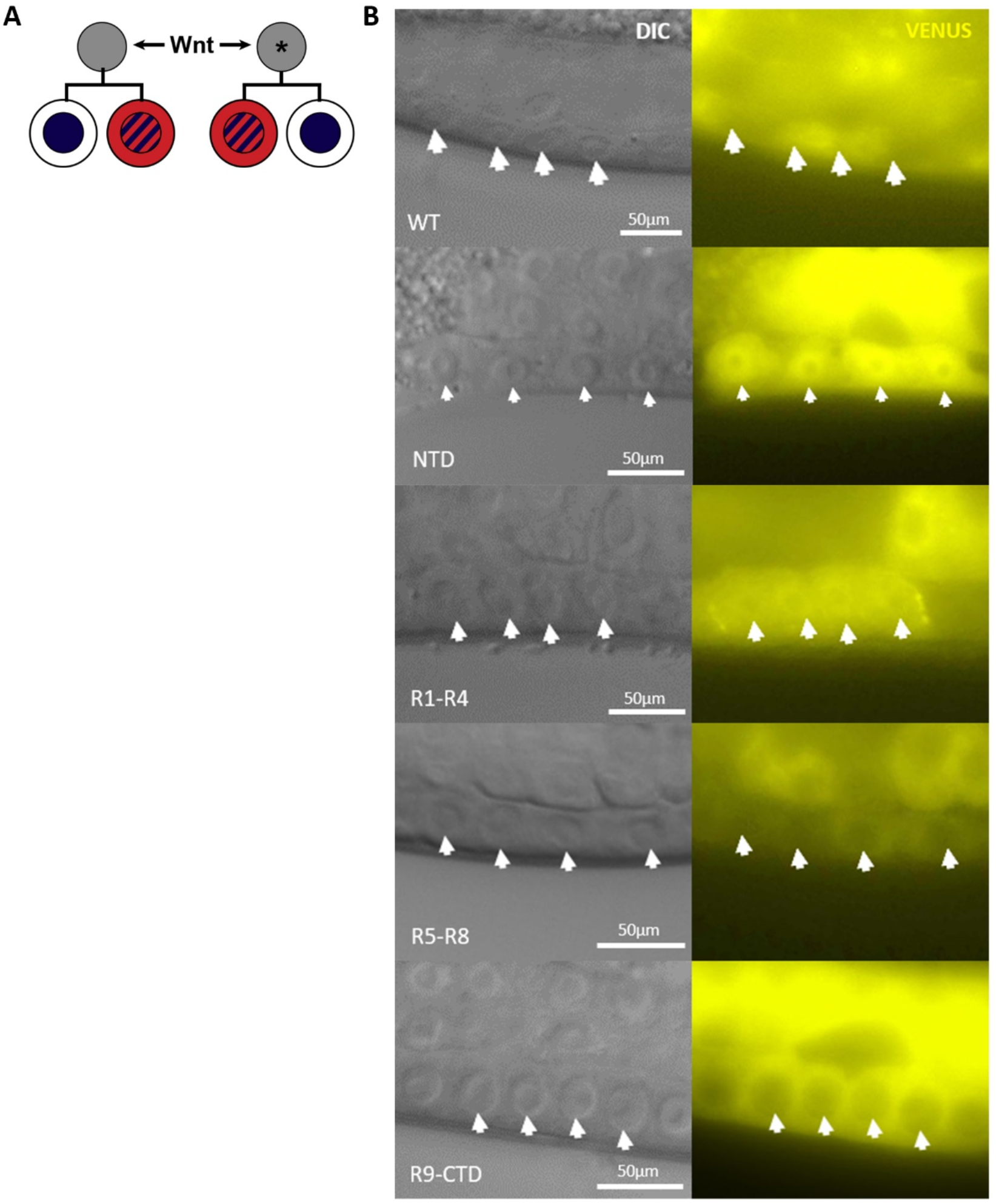
SYS-1 NTD and R1-R4 localizes in the nucleus of VPC. (A) WβA signaling in vulva. Wnt source polarizes two mother VPCs producing an asymmetric set of daughters with a high and low SYS-1 in the nucleus. Asterisk indicates the modified pattern of ACD polarity seen in the posterior vulva. (B) DIC (left) and YFP (right) images of full length SYS-1 and SYS-1 truncations fused to VENUS in vulva. Arrows denote the four central cells in the VPC lineage that are the products of two ACDs.

We tested the nuclear localization of four different SYS-1 truncations, each consisting of roughly 25% of the full-length protein, in the VPCs. The truncations selected for this experiment were the unstructured NTD, R1-R4, R5-R8 (the known interaction site of SYS-1/POP-1) and R9-CTD. We found that, while full length SYS-1 localizes asymmetrically in the nucleus of the proximal daughters and is not detectable in the distal daughter cells (Figure 4B, top), all of the above SYS-1 truncations resulted in symmetric localization between the VPC daughter pairs (Figure 4B). The loss of asymmetric regulation indicates that cytoplasmic regulation of SYS-1 involved in ACD is compromised and it is thus likely that multiple sites spanning more than one of our chosen fragments within SYS-1 is required for proper SYS-1 asymmetric localization. Interestingly, we found that NTD and R1-R4 are both enriched in the nucleus while R5-R8 and R9-R12 are excluded from the nucleus (Figure 4B). This nuclear localization pattern of the NTD and R1-R4 domains is consistent with the presence of two cryptic nuclear localization signals that were not identifiable based on sequence analysis alone, which led us to investigate the mechanism controlling the SYS-1 subcellular localization.

### XPO-1 asymmetrically regulates the nuclear level of SYS-1

To identify regulators of SYS-1 nuclear localization we sought to establish a catalog of SYS-1 physical interactors. To date, the only known direct interactor of SYS-1 is POP-1 (Liu et al., 2008) prompting our use of a mass spectrometry screen to identify novel SYS-1 interactors. We identified several SYS-1 interactors of interest including proteins associated with proteasomal degradation pathway (RPN-5, RPN-9, RPN-10, UBA-1), chromatin remodeling proteins (NAP-1, CEY-2), and nuclear import/export proteins (nuclear exporters XPO-1 and XPO-2 and the importin β IMB-3). We tested the role of the XPO1/CRM1 homolog XPO-1 in SYS-1 nuclear transport. Within mammalian systems, XPO1/CRM1, the ortholog of XPO-1, is a major nuclear exporter that binds to leucine rich nuclear export sequences (NES) (Gorlich and Kutay, 1999; Kau et al., 2004). We thus hypothesized that XPO-1 interacts with SYS-1 to regulate SYS-1 nuclear:cytoplasmic distribution and tested this idea using two different assays. We first examined cell fate changes after altering XPO-1 function, since the degree of SYS-1 nuclear localization would be affected in its function as a POP-1 co-activator, and by directly analyzing SYS-1 localization levels using Venus tagged to full-length SYS-1.

We again utilized the SGP lineage to assay cell fate changes when XPO-1 was depleted in larvae. Since SGPs divide asymmetrically to give rise to DTC, DTC counts were performed at the L3 stage following *xpo-1* RNAi. Surprisingly, DTC quantitation of worms exposed to *xpo-1* RNAi targeting the SGP lineage tissue showed no significant changes in DTC specification (2.7% Sys, n=165) compared to empty vector (2% Sys, n=324). Negative results from the SGP *xpo-1* RNAi could be attributed to protein stability of XPO-1 and its inheritance from previous cell divisions. To circumvent the persistence of inherited XPO-1, we treated worms with Leptomycin B (LMB, 2ng/mL (Perander et al., 2001; Rubio-Solsona et al., 2018)) to block XPO-1 function. Synchronized L1 worms were grown in liquid culture containing either OP50 bacteria or OP50 supplemented with LMB and DTCs were quantified at the late L3 stage. This treatment proved effective in altering DTC number, however, instead of our expected phenotype of DTC increase, LMB treatment decreased DTC count (~23.3% Sys, n=163) when compared to control worms lacking LMB (~1.1% Sys, n=136, p<0.005). The loss of DTC from LMB treatment could be attributed to a confounding role of XPO-1 regulation of POP-1 nuclear export, which is essential for the signaled fate. XPO-1 is a major exportin in worms and have been shown to interact with POP-1 and WRM-1 as well as regulate POP-1 and WRM-1 nuclear asymmetry (Lo et al., 2004; Nakamura et al., 2005). Since this role of XPO-1 may complicate the analysis of SYS-1-dependent cell fate, we therefore turned to direct measurements of SYS-1 nuclear localization to evaluate the role of XPO-1.

Since L1 feeding of *xpo-1* RNAi appears highly deleterious to progression to L3 stage, we instead analyzed the nucleo-cytoplasmic distribution of SYS-1 in EMS daughters. To differentiate the phenotype resulting from WRM-1 and POP-1 loss of asymmetry and SYS-1 loss of asymmetry, we observed SYS-1 nuclear localization in the EMS daughters in the embryos. Embryos were harvested from RNAi-treated adults to obtain a population of embryos that have a varying degree of RNAi penetrance from normal developing embryos as well as embryos with severe developmental delays. We found that among the embryos observed capable of progressing through EMS division (n=6), E and MS daughters exhibit various degrees of SYS-1 nuclear localization (Figure 5A,B), likely due to degree of RNAi efficacy. However, consistent with a role in nuclear export in both daughters, XPO-1 depletion caused an average elevation of SYS-1 nuclear levels in both E and MS compared to embryos from worms subjected to control RNAi. Interestingly, the effects of XPO-1 loss were more severely seen in the MS nucleus than the E nucleus, causing a near loss of asymmetry between the two daughters of EMS. These data indicate that both E and MS utilize XPO-1 to facilitate SYS-1 nuclear export and suggest XPO-1 has a greater role in SYS-1 export from the MS nucleus compared to E (Figure 5B).

**Figure 5.**
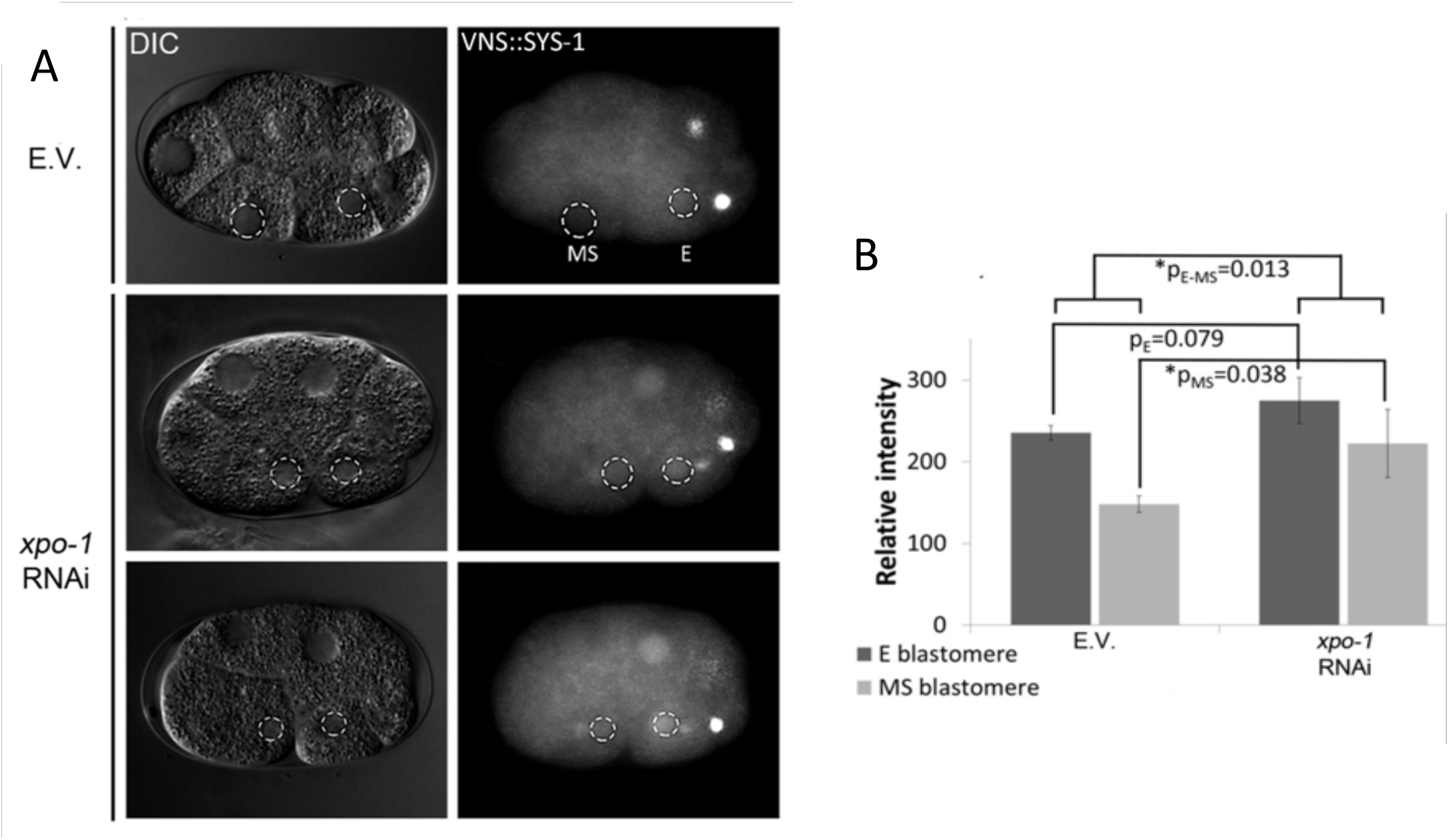
*xpo-1* regulates nuclear SYS-1 asymmetry. (A) *xpo-1* depletion causes a loss of SYS-1 nuclear asymmetry. Nuclear localization of VENUS::SYS-1 following empty vector or *xpo-1*(RNAi) and (B) quantification on embryos observed showing decreased E/MS asymmetry after *xpo-1*(RNAi) (n=6) (right).

## Discussion

Here, we investigate the molecular evolution and the transactivation function of SYS-1, a key regulator in the WβA pathway. We show that SYS-1 is the result of divergent evolution after serial β-catenin gene duplications and sub/neofunctionalizations in nematode evolution. We find that, while the conserved transactivation site in β-catenin is often located towards to carboxyl-terminus of the protein, the transactivation domain of SYS-1 is located towards to amino-terminus of SYS-1 as shown in yeast one-hybrid and *C. elegans* Wnt-induced cell fate assays. Within the signaled daughter of the WβA pathway, stabilized SYS-1 translocates into the nucleus where it interacts with POP-1 and activates transcription of target genes (Kidd et al., 2005; Phillips et al., 2007). Our data support the model that a minimal transactivator domain (NTD and R1-R4) allows of activation of transcriptional target genes once in close proximity with co-activators. Furthermore, as minimal transactivator, each of these fragments by itself is shown to change cell fate specification. From an interactor screen, we found an importin-β-like protein, IMB-3, and an exportin, XPO-1 as candidates for SYS-1 nuclear import and export. XPO-1 is required to prevent aberrantly high nuclear SYS-1 after ACD and for promoting nuclear asymmetry between daughter cells. Localization experiments using tagged SYS-1 fragments uncovered a new cryptic NLS within the N-terminus half of SYS-1 (NTD and R1-R4) that facilitates SYS-1 nuclear localization. Our analyses thus support a model whereby SYS-1, through NLS domains located in the NTD and R1-4, is preferentially imported into the nucleus of the Wnt signaled daughter cell. In the nucleus SYS-1 interacts with POP-1 via the central armadillo repeats and activates transcription of Wnt target genes via two distinct transactivation domains in the NTD and R1-4 (Figure 6, left). In the Wnt unsignaled daughter, SYS-1 is degraded by a destruction complex that includes APR-1/APC and KIN-19/CKIa (Baldwin et al., 2014) and possibly preferentially exported from the nucleus (or experiences a relative lack of nuclear import), resulting in a decrease in Wnt target gene activation (Figure 6, right).

**Figure 6.**
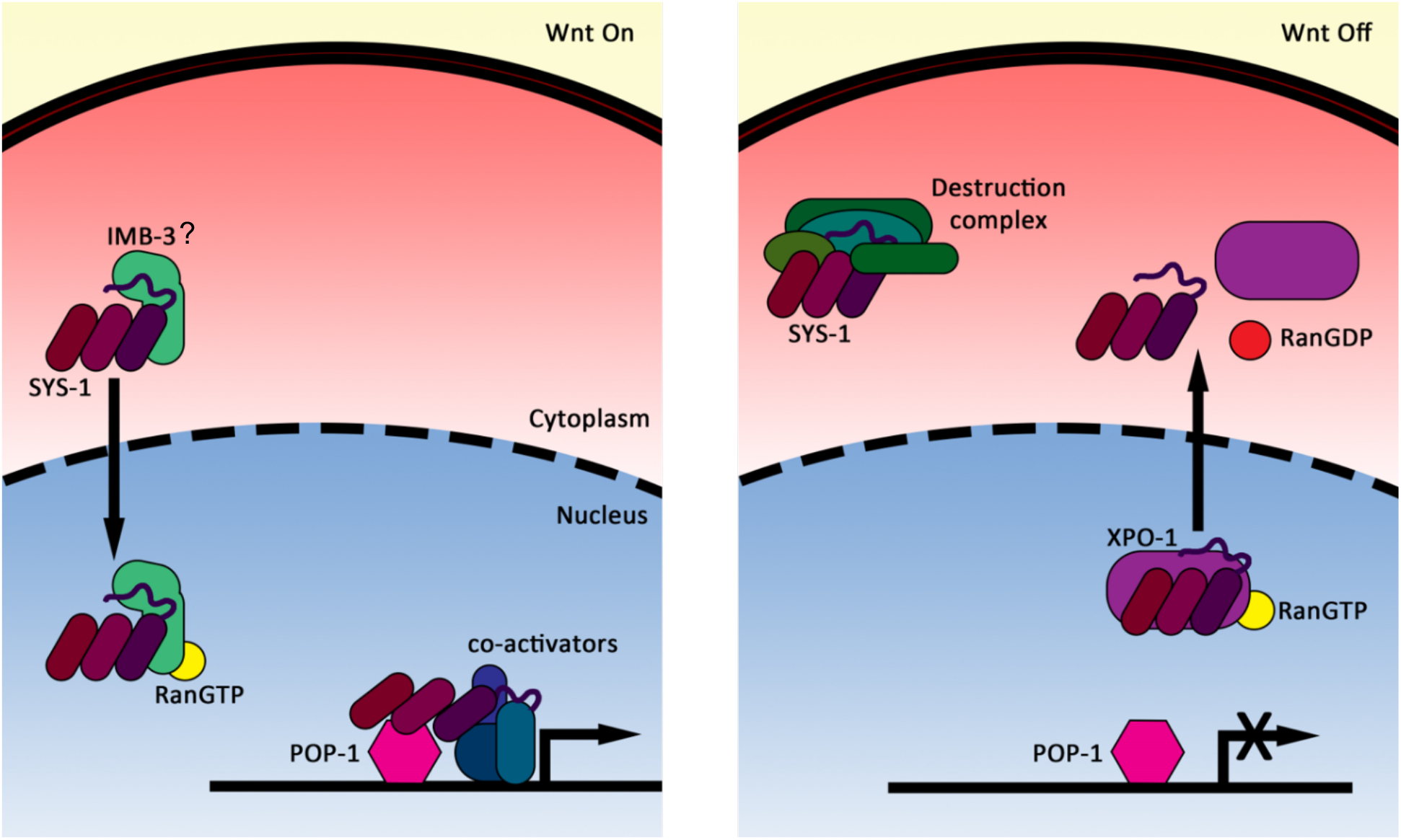
Model of SYS-1 nuclear transport and transactivation. (Left) In the signaled daughter, SYS-1 is imported into the nucleus through the interactions between two of SYS-1 nuclear localization domains located in the NTD-R4, possibly via importer IMB-3. The complex is transported into the nucleus where SYS-1 disassociates with IMB-3 and binds POP-1/TCF via the R5-8 domain. SYS-1 mediates POP-1 interaction with co-activators via SYS-1 NTD and R1-R4. (Right) In the unsignaled daughter, the export of residual SYS-1 in the nucleus is assisted by XPO-1. In the cytoplasm, the destruction complex (e.g. APR-1/APC) binds to multiple sites along the length of SYS-1 to target SYS-1 for proteasomal degradation.

Since mammalian β-catenin has previously been shown to be able to bind to itself (Cox et al., 1999), one intriguing possibility is that β-catenin truncations alter self-regulation. One of the earliest experiments to suggest β-catenin self-interaction was a yeast two-hybrid assay of the CTD of fly *arm* against various truncations of armadillo repeats. This assay showed the strongest binding affinity to the CTD was the central Arm repeat region (Cox et al., 1999), demonstrating a role in β-catenin self-inhibition with respect to CTD interactors. Additionally, the CTD and NTD promote β-catenin stability by shielding the central armadillo repeats from interacting with the destruction complex and a β-catenin truncation that lacks a CTD was degraded faster than full-length β-catenin (Mo et al., 2009), suggesting the CTD promotes β-catenin stability. Since SYS-1 loses the unstructured CTD during its molecular evolution, the loss of the CTD could abolish its self-shielding ability and result in an inherently less stable protein. This is consistent with the elevated turnover of SYS-1 expected to be required in embryonic ACDs that utilize reiterative stabilization and degradation cycles of SYS-1 in as little as 15 minutes. Alternatively, the self-stabilizing role may have shifted to the NTD and represent a novel mechanism to stabilize SYS-1 by binding to the armadillo repeat containing domain.

The NTD as well as the CTD of β-catenin are important for proper Wnt signaling in mammals and Drosophila (Cox et al., 1999; Valenta et al., 2011). For example, in *D. Melanogaster*, truncation of the CTD decreases transcriptional activity compared to wild type (Cox et al., 1999). Among co-activators and interactors that interact with β-catenin includes TCF in the central armadillo repeats and BCL-9 in the NTD, which is essential in recruiting Pygopus (Liu et al., 2008; Mosimann et al., 2009; Valenta et al., 2011) leading to transcriptional activation. In addition to co-activators, β-catenin also binds to a host of chromatin-modifying factors; histones acetyltransferases such as CBP, SWI/SNF factor BRG-1 and the mediator component MED12 in its CTD region (Barker et al., 2001; Hecht et al., 2000; Kim et al., 2006). We find that SYS-1 R5-R8 and R5-CTD were excluded from the nucleus yet nevertheless influence SGP cell fate specification (loss of DTC) in worms containing endogenous SYS-1 (Figure 3B). One possible explanation for the loss of DTC could be that these SYS-1 fragments are binding POP-1 and therefore preventing the POP-1/SYS-1 interactions in the nucleus. In this case, POP-1/R5-R8 or POP-1/R5-CTD would be shuttled into the nucleus via the POP-1 NLS and therefore prevent endogenous SYS-1 from binding POP-1. The caveat to this hypothesis is that a loss of DTC would necessarily be attributed to a high amount of POP-1/R5-R8 being trafficked into the nucleus. However, we do not observe such nuclear localization of our C terminal SYS-1 fragments. While it remains possible that a relatively small amount of SYS-1 fragments could effectively sequester sufficient amounts of POP-1 to decrease Wnt target gene activation, current models predict that a decrease of up to 50% in unbound POP-1 results in activation of target genes, rather than repression. Thus, this model would have to invoke very little free POP-1 in the nucleus in order to sufficiently impair Wnt target gene activation. Given these observations, the POP-1 sequestration model seems unlikely, though it cannot be ruled out at this time.

Alternatively, we propose that R5-R8 and R9-CTD could have a self-regulating function that is remnant of SYS-1 molecular evolutionary past and the truncation prevents transcription of target genes by preventing SYS-1 from entering the nucleus through the interaction with endogenous SYS-1. In this scenario, the loss of DTC could be attributed to R5-R8 and R9-CTD interactions with endogenous SYS-1 NTD and R1-R4, masking SYS-1 NLS and therefore preventing proper SYS-1 trafficking. This model could better explain the exclusion of R5-R8 and R9-CTD from the nucleus and the loss of DTC in transgenics containing the above-mentioned truncations.

Canonically, Wnt signaling results in β-catenin stabilization and nuclear translocation. Various models have been proposed for β-catenin nuclear import within metazoans and a definitive reconciling model has yet to be developed. One of the earliest studies in this field proposes that β-catenin nuclear import was independent of importins. In this model, β-catenin moves across the nuclear membrane due to direct interaction with the nuclear pore complex through interaction with nucleoporin FxFG repeats, rather than interactions with importins, due to structural similarities between the armadillo repeats found in β-catenin and the HEAT repeats found in importin-β. (Fagotto et al., 1998; Yokoya et al., 1999). Alternatively, Lgs/BCL9 has been suggested to escort beta-catenin into the nucleus where both proteins are anchored through interactions with Pygopus (Townsley et al., 2004). More recently, a role for importins was identified in glioma cells, where depletion of importin-11 resulted in a cytosolic shift in the nuclear-cytoplasmic beta-catenin distribution, suggesting importins could regulate nuclear β-catenin levels during tumorigenesis (Ni et al., 2021). Similarly, we report a SYS-1 interaction with importin β IMB-3 suggesting the hypothesis that conserved importin interactions may lead to SYS-1 nuclear translocation.

Subsequent studies suggested an APC role in β-catenin export (Henderson, 2000). Similarly, β-catenin nuclear export depends on passive transport via the HEAT tandem motif in the armadillo repeats or through active export through the interactions with CRM1 via APC. While the previous study showed evidence that suggest that APC is required for CRM1 mediated nuclear export of β-catenin (Henderson, 2000; Henderson and Fagotto, 2002), other studies suggest an APC-independent model of β-catenin export (Eleftheriou et al., 2001). For instance, β-catenin rapid export was shown to be independent of APC and CRM1 although overexpression of CRM1 will moderately stimulate an export of β-catenin. Here, we report an interaction between SYS-1 and XPO-1/CRM1 and a requirement for XPO-1 in lowering nuclear SYS-1 levels, especially in the unsignaled daughter. Our model (Figure 6), whereby the daughters of a Wnt-regulated ACD exhibit differential cytoplasmic and nuclear SYS-1 levels, suggests multiple mechanisms to differentially control SYS-1 concentration in different subcellular locales that reinforce each other to drive proper SYS-1 levels, leading to differential transcriptional status in daughter cells and subsequent cell fate asymmetry.

## Materials and Methods

### Plasmid generation

Cloning was carried out using Pfu Turbo polymerase. All plasmids transformed into ONESHOT™ TOP10 chemically competent E. coli cells (Catalog number: C404006) unless mentioned otherwise. Transformed cells were sequenced prior storage in −80°C for future use. All plasmid generated for the yeast-one hybrid assay used pBTMKnDB as a backbone vector.

### Strain generation and maintenance

Strains were maintained using standard *C. elegans* methods [Brenner, 1978]. Refer to Supplemental Table 1 for list of worm strains used. Transgenics were created through coinjecting truncated plasmids with either Pttx-3:GFP (for PHS:mCherry::SYS-1trunc) or Pttx-3:mCherry (for PSYS-1:VENUS::SYS-1trunc) co-injection marker with a corresponding plasmid in a 1:1 ratio. Transgenic carrying arrays of truncations were maintain by selecting for positive co-injection marker. Transgenics used in chapters below were selected from multiple lines created during co-injection based on the array inheritance, the ease of maintenance, and a strong expression pattern. Stock strains were obtained from Caenorhabditis elegans Genetics Center unless otherwise noted.

### Yeast one-hybrid assay and yeast maintenance

Yeast strains were maintained using standard methods. ONPG assay were performed using an adapted protocol from Reynolds et al (Reynolds et al., 2001).

### Microscopy

Transgenics were synchronized at L1 and plated on OP50 lawn on NGM for approximately 22 hours in 25°C. Larval worms were then immobilized using 10μM muscimol on a 5% agarose gel pads. Embryos were obtained from dissecting adult C. elegans in M9 buffer before mounted on 3% agarose gel pads. Cells and their nuclei were identified using differential interference contract (DIC) before fluorescence imaging.

### RNAi treatments

RNAi on specific target were done through feeding worms bacteria expressing dsRNA. For negative RNAi controls, worms were fed bacteria containing an empty vector (L4440). For the purposes of DTC count, worms were synchronized at L1 before plating on RNAi specific plates until late 3rd larval stage. For EMS imaging, worms were synchronized and plated on OP50 NGM plate until late adult. Worms were then washed and transferred to RNAi plates. Embryos were dissected from mothers for imaging. Duration of RNAi varied based on the specific gene.

### Distal Tip Cell counts

Transgenic *P_lag-2_::GFP* larvae were synchronized at L1 and plated on OP50 lawn on NGM for until late L3 stage for DTC counts via GFP visualization. % Sys was calculated based on the percentage of SGP that failed to produce a DTC.

### Mass spectrometry screen

Both N2 Bristol (wild type) and BTP50 (PHS:mCH::SYS-1::POP-1DBD) were maintained in liquid cultures and adult worms were isolated via a sucrose gradient protocol. Worms were then mechanically lysed using the bead beater homogenizer. Immunoprecipitation protocol using RFP antibody (a kind gift from lab of Thomas Rutkowski at the University of Iowa) targeting mCherry were performed on the lysates. Samples ran on NuPAGE™ 4-12% Bis-Tris Protein Gels and subsequently Coomassie stained. Bands appearing on BTP50 but absent on N2 were excised and sent to University of Iowa Proteomics Facility for protein identification via tand mass spectrometry using a Bruker UltrafleXtreme MALDI TOF/TOF Mass Spectrometer

### Leptomycin B treatment

Worms were synchronized at L1 and were incubated in M9 or M9 and 2ng/mL LMB for 2.5 hours. Worms were washed with M9 twice and plated on OP50 lawn on NGM plates. LMB were diluted in 95% ethanol and similar concentration of ethanol were added into control group to compensate for the presence of ethanol in LMB group.

## Acknowledgements

We thank members of the Phillips lab for helpful comments on the manuscript. Some strains were provided by the Caenorhabditis Genetics Center, which is funded by the National Institute of Health (NIH) Office of Research Infrastructure Programs [P40 OD01440]. This work was supported by NIH GM114007 (BTP) and the Ballard-Seashore Dissertation Fellowship (AKW). We thank the laboratory of Thomas Rutkowski (UI) for the anti-RFP antibody used in immunoprecipitations.

