## Supplemental Materials for "Nuclear localization and transactivation of SYS-1/β-catenin is the result of serial gene duplications and subfunctionalizations"

**Supplemental Figures**

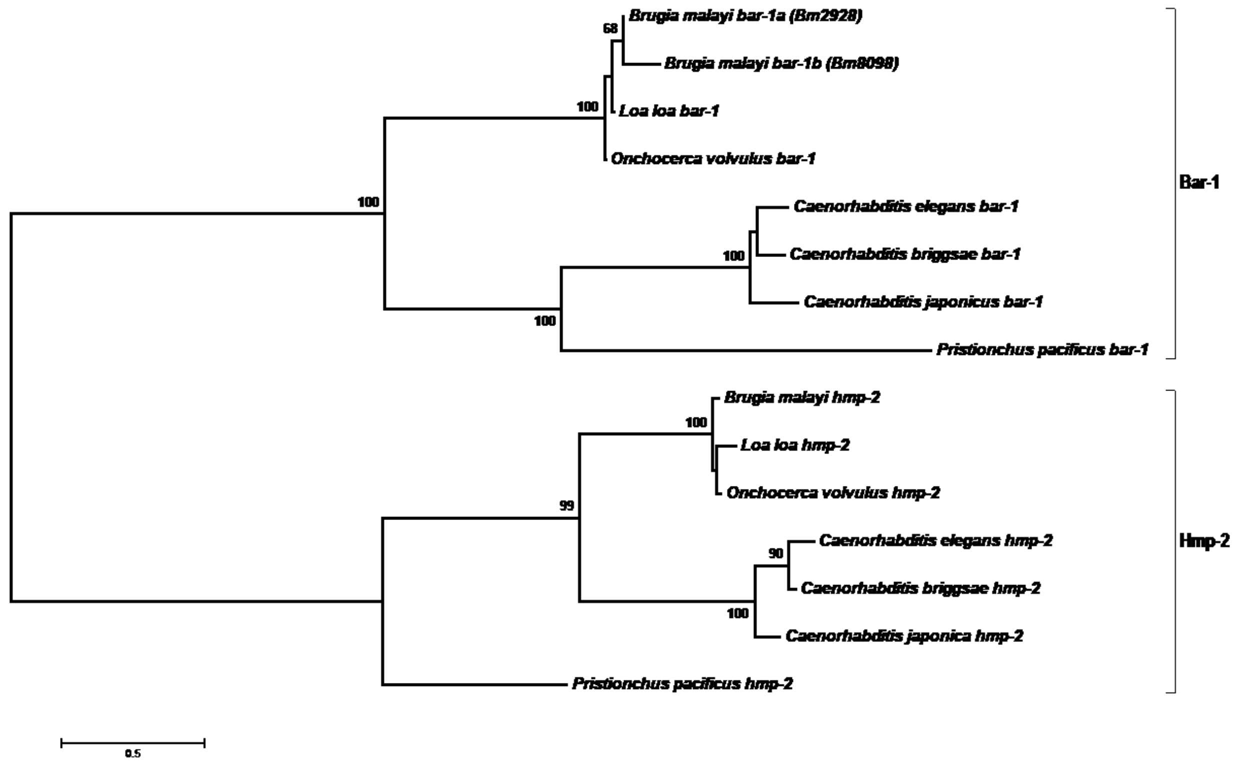


Supp. Figure 1 **The nematode genes *hmp-2* and *bar-1* represent sub-functionalized duplications of armadillo in the latest common ancestor of all nematodes.** All nematodes maintain the two sub-functionalized armadillo genes *bar-1* and *hmp-2*, which have separately maintained roles in transcription and adhesion, respectively. Note that both gene clades have comparable amounts of divergence since the latest common ancestor of nematodes. Shown is the optimal phylogenetic tree which was computed using the Neighbor-Joining method (Saitou and Nei, 1987). The percentage out of 500 bootstrap replicate trees in which the associated taxa clustered together is shown next to the branches (Felsenstein, 1985). The evolutionary distances were computed using the JTT matrix-based method (Jones et al., 1992) and are in the units of the number of amino acid substitutions per site. The rate variation among sites was modeled with a gamma distribution (shape parameter = 1). All positions with less than 55% site coverage were eliminated. That is, fewer than 45% alignment gaps, missing data, and ambiguous bases were allowed at any position. There was a total of 692 positions in the final dataset. All NJ analyses in this study were conducted in MEGA6 (Tamura et al., 2013).

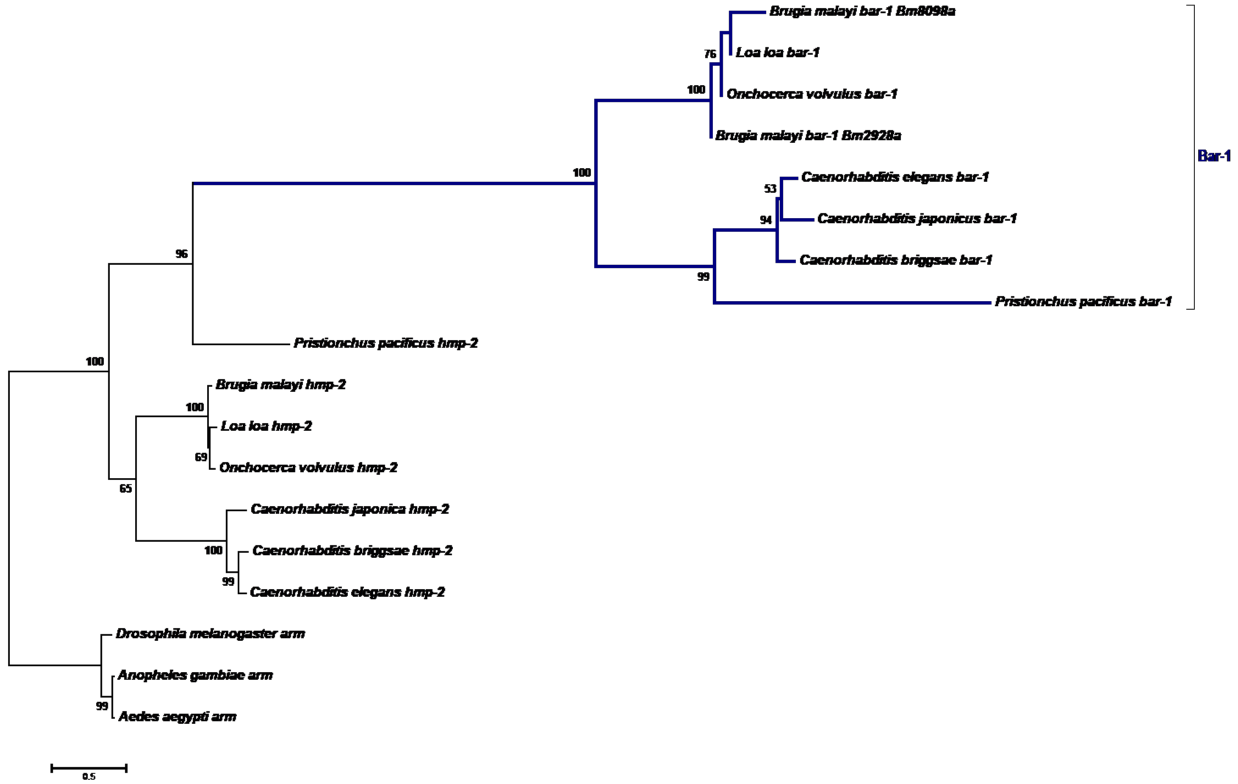


Supp. Figure 2 **The nematode armadillo paralog *bar-1* is neofunctionalized relative to the hmp-2 paralog.** Including other ecdysozoan outgroup sequences from flies, which have not been duplicated, shows that after the nematode duplication there was rapid evolution in the *bar-1* locus prior to diversification of extant nematodes, suggesting either neofunctionalization or at last fast rate of evolution of bar-1. Shown is the optimal phylogenetic tree computed using the NJ method [1]. The percentage out of 500 bootstrap replicate trees in which the associated taxa clustered together is shown next to the branches [2]. The evolutionary distances were computed using the JTT matrix-based method [3] and are in the units of the number of amino acid substitutions per site. The rate variation among sites was modeled with a gamma distribution (shape parameter = 1). All positions with less than 40% site coverage were eliminated. That is, fewer than 60% alignment gaps, missing data, and ambiguous bases were allowed at any position. There were a total of 838 positions in the final dataset.

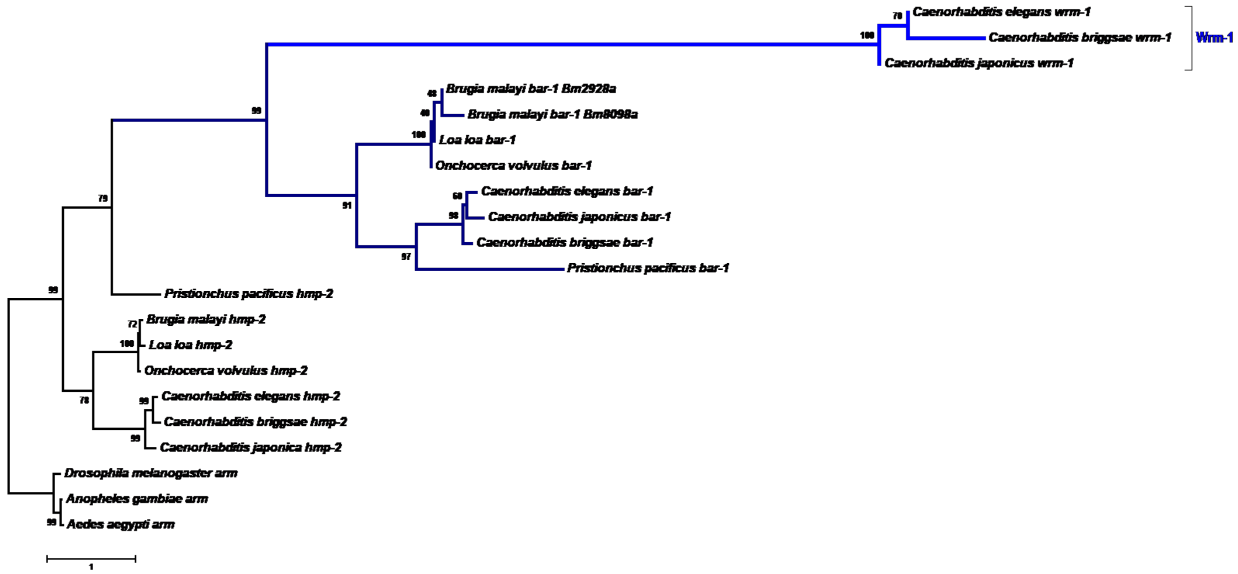


Supp. Figure 3 **The gene *wrm-1* is a more recently neo-functionalized version of nematode *bar-1*.** The *wrm-1* gene is a more recent neofunctionalization of *bar-1*, much like *bar-1* was a neofunctionalized duplication of nematode arm. Accordingly, the amount of sequence alignment and evidence of homology of *wrm-1* to the arm family was doubly reduced by these events. Shown is the optimal phylogenetic tree computed using the NJ method [1]. The percentage out of 500 bootstrap replicate trees in which the associated taxa clustered together is shown next to the branches [2]. The evolutionary distances were computed using the JTT matrix-based method [3] and are in the units of the number of amino acid substitutions per site. The rate variation among sites was modeled with a gamma distribution (shape parameter = 1). All positions with less than 40% site coverage were eliminated. That is, fewer than 60% alignment gaps, missing data, and ambiguous bases were allowed at any position. There were a total of 826 positions in the final dataset.

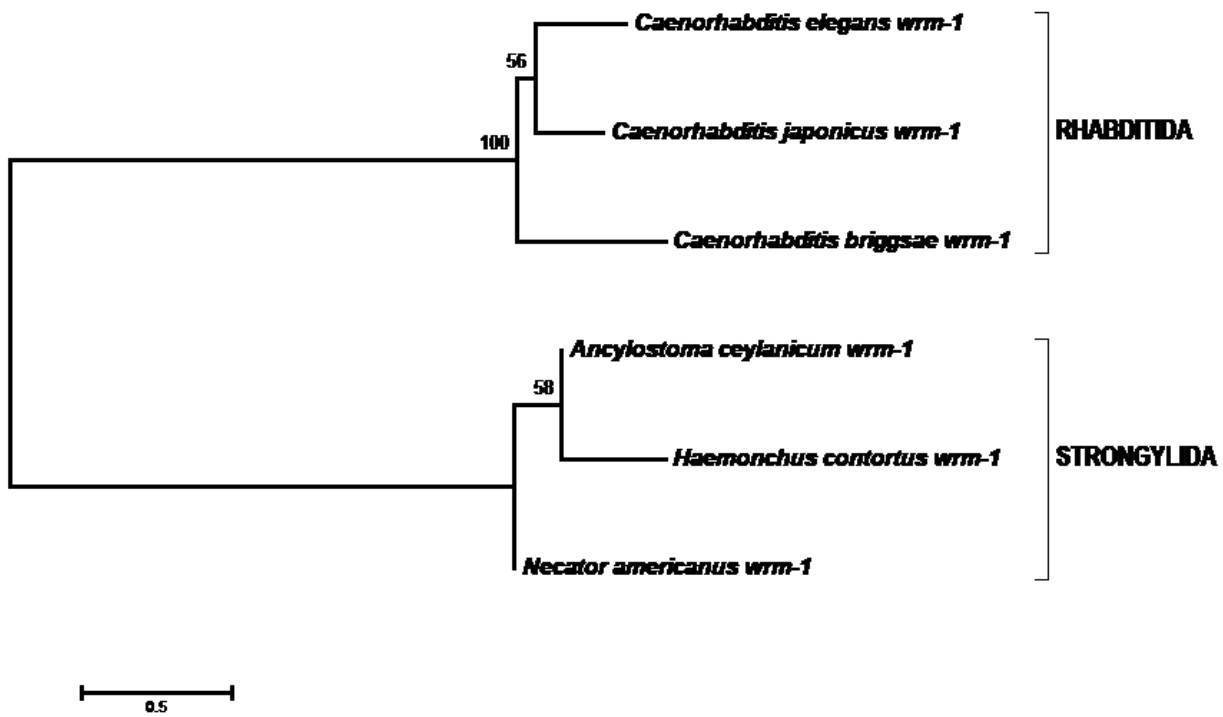


Supp. Figure 4 **The *wrm-1* gene originated by the time of the latest common ancestor of the sister nematode orders RHABDITIDA and STRONGYLIDA.** The gene *wrm*-1 is also present in the two sister nematode orders of RHABDITIDA and STRONGYLIDA but not in any other nematode order. Shown is the optimal phylogenetic tree computed using the NJ method [1]. The percentage out of 500 bootstrap replicate trees in which the associated taxa clustered together is shown next to the branches [2]. The evolutionary distances were computed using the JTT matrix-based method [3] and are in the units of the number of amino acid substitutions per site. The rate variation among sites was modeled with a gamma distribution (shape parameter = 1). All positions with less than 50% site coverage were eliminated. That is, fewer than 50% alignment gaps, missing data, and ambiguous bases were allowed at any position. There were a total of 785 positions in the final dataset. Evolutionary analyses were conducted in MEGA6 [4].

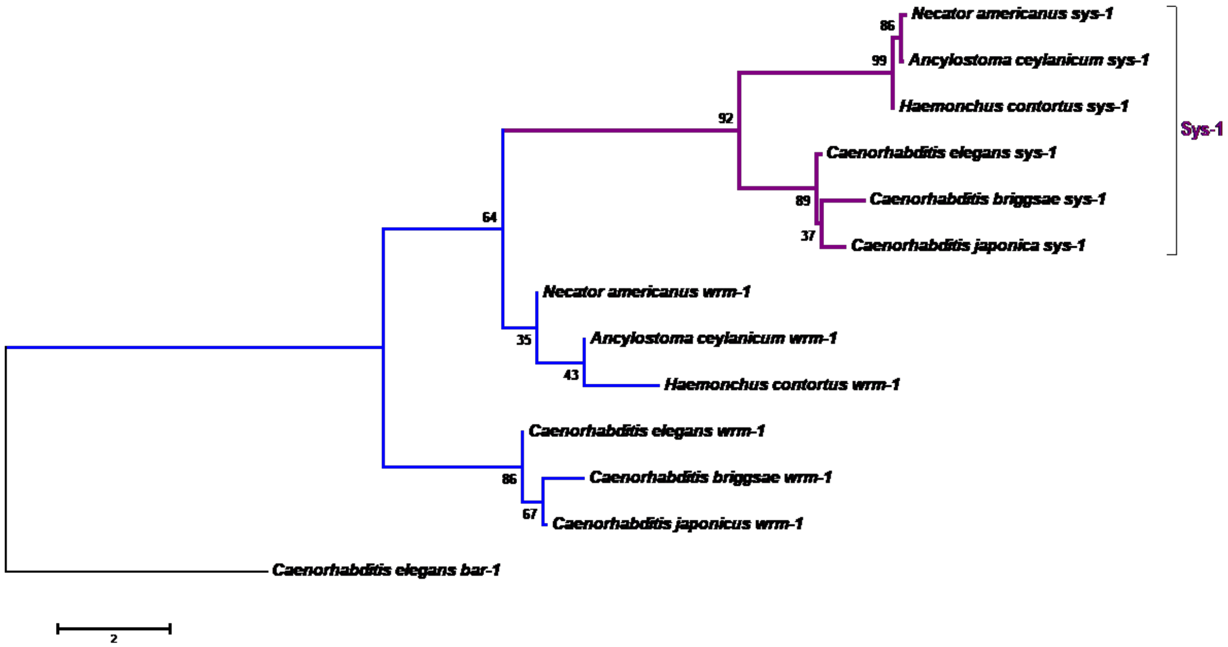


Supp. Figure 5 **The gene *sys-1* is a neo-functionalized duplication of the doubly-neofunctionalized *wrm-1.*** Shown is the optimal phylogenetic tree computed using the NJ method [1]. The percentage out of 500 bootstrap replicate trees in which the associated taxa clustered together is shown next to the branches [2]. The evolutionary distances were computed using the JTT matrix-based method [3] and are in the units of the number of amino acid substitutions per site. The rate variation among sites was modeled with a gamma distribution (shape parameter = 1). All positions with less than 45% site coverage were eliminated. That is, fewer than 55% alignment gaps, missing data, and ambiguous bases were allowed at any position. There were a total of 800 positions in the final dataset. Evolutionary analyses were conducted in MEGA6 [4].

```
Hc wrm-1:  MPRNVFCHFDQSPLRSHQLGLTVNAFMDPMSSAYAVPSPYDQQVPSTSSPSCAMARPNTPQTPYG--RSAFGSAHASPFRPANAAD
Hc sys-1:  MDAKQVY-DQQIPSTSQFSHYPKPEMMQQYPPACRQNMQHMYPASYPRSPSRVPQTDHTSHWLSGPYNAVYANSPVPSGAPSVMSH

AVRLRETCAARNAQTAQWMDSQYLAGVSGYHGSGVQSGPCSVMSAASGGGMSMMSGMSQMSLGNMSQLSICTQLTEAHYTQNAAYQTTSGASSVENI
VSMQDDLVSVMSVQTTLNASTAPCPQDSQLTELRPQSQFSSSASTPP--CTENVDPATLQEVRRNIAYSINNFYSNEQFKIIQCVCHYAQRNMLQFV

DPSNIFEINRQVNTLIMSLRHEDPEIRRKAANSVQSMARQRLLAMVSNHATLRELICHFLQFVLEHTIEQNITQLMQSIFNILNCRTCFVVMQDRYI
DMGSVHKLLLLITQLLKNT--IDPNFRRQNSLQTSNLLLNTIFYLSRNQTTLR-VVMHVASDKNTSEFIYIIAHFAKVNLNWAAALTLHTLFDYNGP

QQILVHIIAKYLMPDHPFTAWTVIAFTAILRKKDLRGFRRMCRNEYFVRQLMIILGSTSSSHKSLIIDSIRMLITRNLPLKEFFVRSKGMEKLLEFM
EGAKCITYAKQIQPG-----AVLDALLHILKEKPDR------KRTLFVLHSLQLLARKDNNAKQHIVQSY------NGRFIELLIGSAQGGLLVLAD

SCEHEEKVLRSSATTLLTLISTESAKYGIEFVKLGGVQIFGNLLSHGSTMLLHEVLHCLAGVGDLKEALETQDISNTIVSVLQLMG---AHDPILVN
NARDQGRLLYRILALLNVVMASQTAAE--LFVRWGGLQKVCERLYQ-STDVLREGIHLIALASDVKTVDD-QDLRVSIRQVLDILCRSNDRDLFVRR

HGTAFLLNVSANSIRNKASMVTAQAPDTLLSVLNHRNNYLTIP-LANVRQLIASITDNVLICLANLTRN--QDECGRAACIQIARKPDSAKTLLNVL
YSSGFLVNVIANRPVNKRLLIECNAFDIFTRLCYQYSNLEQWSQTKEMRNMLEELVDNILLCLNSLIVEPLKPDVTKARDTCLSQKS-FQVCLLKLL

ASGTVECQNRVLMVIHSTCICVAEYKLFFLDTVDDYNQSNLVTFTLRLLWNNFVQYNSFSEAERIHRRKQIDRVLRLLATLSKSEELLQQITAIFRE
SNGSQCIRARVLRLMTLLLSCCAGWGATVTQAVLEDKGENIIGVVFNVIS---ITRADATDDNRRETVNILTACFRVLNLVMFCDEGAKKTVTLFAR

HNPLPLLRPFRTNDFTAHRDFLEFLDILSSNPMRFDLIPKWLSTP---ESQQILHYYCQFQPDPKLMGHARNIGSRLQQAN-------VEGMQTT-*
NSPVSLMELLRT--FTEDNLVVQILETCDRVADDENLAQLWASDRPLIDANANSSHPEIARAAQSLVSKLSVVNSDFEPLAGVFASMGTEGFSMEM*
```

Supp. Figure 6 **Alignment of the β-catenin proteins encoded by *wrm-1* and *sys-1* of *Haemonchus contortus.*** Homology of the predicted full-length peptide sequences encoded by *wrm-1* (top sequence) and *sys-1* (bottom sequence) of *H. contortus* is demonstrated by the low but homogenous alignment of identical residues (yellow highlighted residues with underlining) throughout the length of the two proteins. Alignment was constructed by ClustalW (Gonnet, gap open = 5.0, gap insertion = 0.05) with a minor adjustment of N- and C-terminal at the candidate start and stop codons.

```
N-terminal domain

H_sapiens          ------------------------------------------------------------     0
M_musculus         ------------------------------------------------------------     0
D_melanogaster     --------------------MS--------------------------------------     2
HMP-2              ------------------------------------------------------------     0
BAR-1              --------------------MDLDPNLVINHD---------------------------D     13
WRM-1              --------------------MDVDCAETFSQPCTPLNFNPMTPSTSRVSTPVRPSSTMSA     40
SYS-1              MHSTGEPQRGPAGPYHHPMPYHVDQIPTTSHPSWHQAPHPGLP-----TPPYHPQQ---H     52

H_sapiens          ---M-----ATQADLM-----------------------EL----DMAMEP-DRKAAVSH     24
M_musculus         ---M-----ATQADLM-----------------------EL----DMAMEP-DRKAAVSH     24
D_rerio            ---M-----ATQSDLM-----------------------EL----EMAMDP-DRKAAVSH     24
D_melanogaster     --YM-----PAQNRTM----------------SHNNQYNPP----DLPPMV-SAKEQTLM     34
HMP-2              ------------------------------------------------------------     0
BAR-1              TNLS-----EASFTME----------------QHTSSYSDI----HMGSTPCTGHRKVDM     48
WRM-1              RQY-SGSPFKAQPQNMEPSNSRVQ---------------------ELREAAVGKRSYTNA     78
SYS-1              PNYISMQPPPSQQQQMTPQALSTQQQQQVQQQQQRQLYSSPSPSRGPAAPAQNQKFRTEQ     112

H_sapiens          WQQQSYL-DSGI-HSGATTTAPSLSGKGNPEEEDVDTSQVLYEWEQGFSQSFTQEQVADI     82
M_musculus         WQQQSYL-DSGI-HSGATTTAPSLSGKGNPEEEDVDTSQVLYEWEQGFSQSFTQEQVADI     82
D_rerio            WQQQSYL-DSGI-HSGATTTAPSLSGKGNPEDDDVD-NQVLYEWEQGFNQSFNQEQVADI     81
D_melanogaster     WQQNSYLGDSGI-HSGAVTQVPSLSGKED---EEMEGDPLMFDLDTGFPQNFTQDQVDDM     90
HMP-2              ----------------------------------------MLLHST--------NSYSIF     12
BAR-1              WRNHNF--DSGF-QTMNHSEAPSIISSLHPSSH-LSGMSSMADYEP--------------     90
WRM-1              WMQGTYAPPQMAGQQQRFSRPPSVIGSTM-------------------------------     107
SYS-1              WINNQAWPYSQ-------SPAPPVAPSVM-------------------------------     134

H_sapiens          DGQYAMTRAQRVRAAMFPETLDEGMQIPSTQFDAAHPTNVQRLAEPSQMLKHAVVNLIN-     141
M_musculus         DGQYAMTRAQRVRAAMFPETLDEGMQIPSTQFDAAHPTNVQRLAEPSQMLKHAVVNLIN-     141
D_rerio            DGQYAMTRAQRVRAAMFPETLDEGMQIPSTQFDSAHPTNVQRLAEPSQMLKHAVVNLINY     141
D_melanogaster     NQQLSQTRSQRVRAAMFPETLEEGIEIPSTQFDPQQPTAVQRLSEPSQMLKHAVVNLINY     150
HMP-2              TDHEVETRTSRIRSAMFPDWIPPTSAAEAT---NSTTSIVEMMQMPTQQLKQSVMDLLTY     69
BAR-1              --------------------------IPTLS-----------------------------     95
WRM-1              -------------------------------------------------SHMTNMSEMTA     118
SYS-1              -------------------------------------------------SYHQ-------     138

H_sapiens          ------------------------------------------------------------     141
M_musculus         ------------------------------------------------------------     141
D_rerio            ------------------------------------------------------------     141
D_melanogaster     ------------------------------------------------------------     150
HMP-2              EGSNDMSGLSLPDLVKLMCDHDESVVARAVHRAYM-----LSREDPNFFNAPGFDHRSFV     124
BAR-1              ------------------------------------------------------------     95
WRM-1              YSYGGLSMLSVNT---EMGEFNNFVNQAPYQRALTRVSQVSENQDP--------------     161
SYS-1              -GDDRMSMLSVNT---TMTN--QFPD---SQRCYSSNGSTCENVNPEMMQM---------     180

H_sapiens          ------------------------------------- 141
M_musculus         ------------------------------------- 141
D_rerio            ------------------------------------- 141
D_melanogaster     ------------------------------------- 150
HMP-2              EALMAASKSSNVNVRRNAIGALSHMSEQRGGPLLIFR 161
BAR-1              ------------------------------------- 95
WRM-1              ------------------------------------- 161
SYS-1              ------------------------------------- 180

C-terminal domain

H_sapiens          DKPQDYKKRLSVELTSSLFRTEPMAWNETADLGLDIGAQ------GEPL-GYRQDDPSY-     52
M_musculus         DKPQDYKKRLSVELTSSLFRTEPMAWNETADLGLDIGAQ------GEAL-GYRQDDPSY-     52
D_rerio            -KPQDYKKRLSVELTSSLFRTEPMTWNETGDLGLDIGAQ------GEPL-GYRQDDPSY-     51
D_melanogaster     -KPQDYKKRLSIELTNSLLREDNNIWANA-DLGMGPDLQDMLGPEEAYEGLYGQGPPSVH     58
HMP-2              ------------------------------------------GDSAAIMNMS--NSYDYE     16
BAR-1              -----YEKREKTRN--TLPRYNSYLESQFGHMSMTTPRSEALNSSGEVCEGA--------     45
WRM-1              ------------------------------------------------------------     0
SYS-1              ----------ENDLLAGIFSNRE-------------------------------------     13

H_sapiens          -----RSFHSGGYGQDALGMDPMMEHEMG--------GHHPGADYPV-------------     86
M_musculus         -----RSFHSGGYGQDALGMDPMMEHEMG--------GHHPGADYPV-------------     86
D_rerio            -----RSFHSGGYGQDAMGMDPMMEHEMA--------GHHPGPDYPV-------------     85
D_melanogaster     SSHGGRAFHQQGYDT--LPIDSMQGLEISSPVGGGGAGGAPGNGGAVGGASGGGGNIGAI     116
HMP-2              MSGSAADWQRDGLERE-------------------------------------------L     33
BAR-1              ----GEQWS------TPLTDDTMMDSYCNS---SGRDSSKPYNS-----------PMYHS     81
WRM-1              ------------------------------------------------------------     0
SYS-1              ------------------------------------------------------------     13

H_sapiens          --DGLPD----------------LGH--A----------------QDLMDGLPPGDSNQL     110
M_musculus         --DGLPD----------------LGH--A----------------QDLMDGLPPGDSNQL     110
D_rerio            --DGLPD----------------LGH--T----------------QDLIDGLPPGDSNQL     109
D_melanogaster     PPSGAPTSPYSMDMDVGEIDAGALNF--D----------------LDAMPTPPNDNNNLA     158
HMP-2              FAEMYPTN-------------------------------DGGHSESINMALNNSQMRPNH     62
BAR-1              PPAMYP--EYSIGPPETYLDPHATASCYPRPTPPQYNSYDRSPPVYNDLPSNPGPSSHSS     139
WRM-1              --------------------QHQMQH--H----------------RQLM-----------     11
SYS-1              ------------------------------------------------------------     13

H_sapiens          AWFDTDL---        117
M_musculus         AWFDTDL---        117
D_rerio            AWFDTDL---        116
D_melanogaster     AWYDTDC---        165
HMP-2              NWYDTDL---        69
BAR-1              DYYPSRNSRF        149
WRM-1              ----------        11
SYS-1              -YF-------        15
```

**Supp. Figure 7 Alignment of the NTD (top) and CTD (bottom) regions of β-catenin protein by various organisms.** Conserved region was found for in *bar-1* (green) or *hmp-2* (yellow) with other organisms but are lacking in other *C. elegans* homologs.

| SYS-1 truncation | Yeast name | Amino acids |
|---|---|---|
| N/A | L40 | N/A |
| Empty vector | pBTMKnDB | N/A |
| NTD-R4 | LA9 | 1-379 |
| R5-CTD | LA10 | 380-811 |
| R3-CTD | L11 | 271-811 |
| R1-R3 | LA14 | 180-379 |
| R3-R4 | LA15 | 217-379 |
| NTD* | LA16 | 1-112 |
| *NTD-R3 | LA26 | 110-331 |
| *NTD-R4 | LA30 | 110-379 |
| Full-length SYS-1 | LA31 | 1-811 |
| NTD | LA54 | 1-179 |
| R1-R4 | LA55 | 180-379 |
| R5-R8 | BP56 | 380-592 |

Supp. Figure 8 List of yeast strains used in Y-1-H assay.

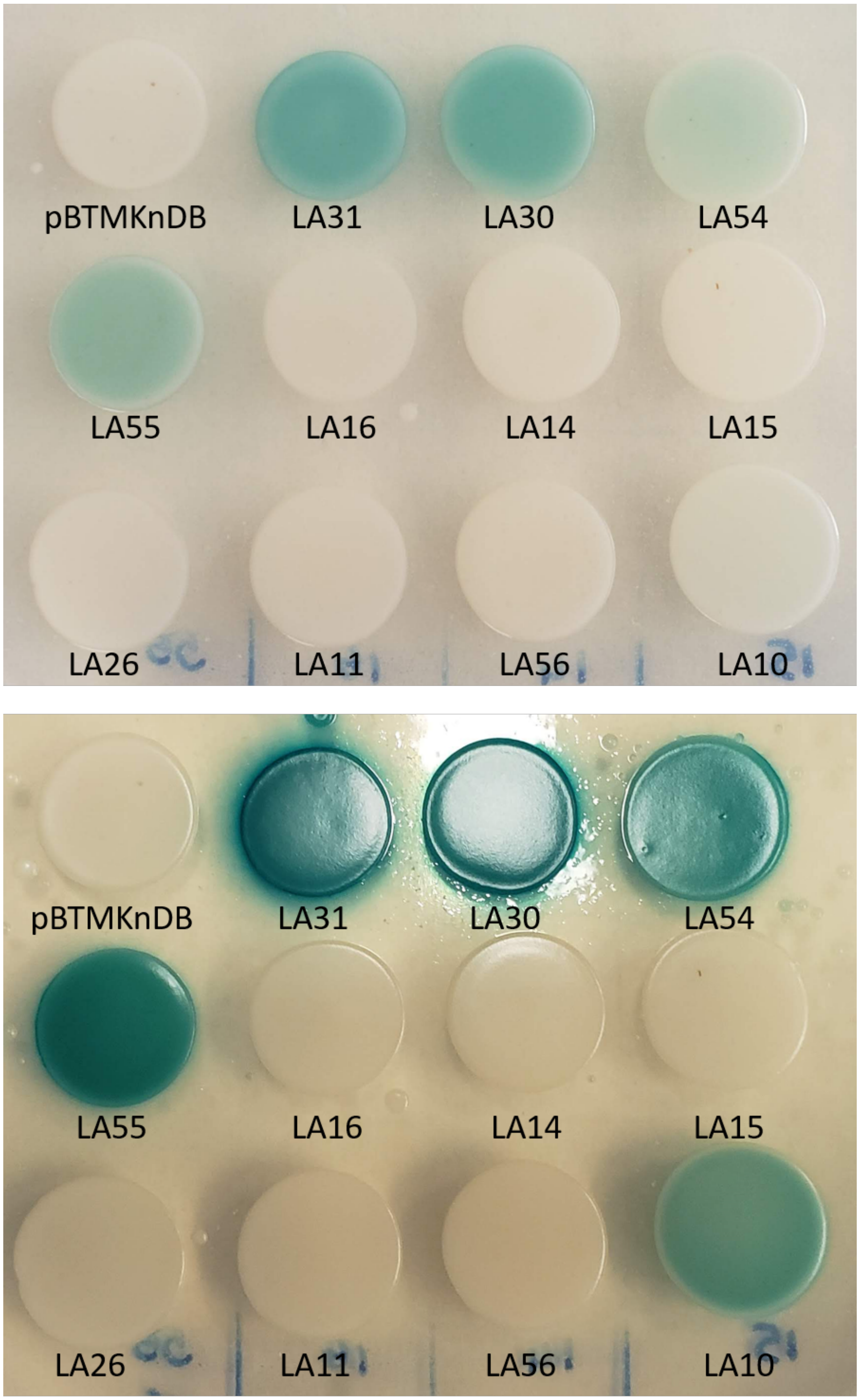


Supp. Figure 9 X-gal assay on Y-1-H constructs. Yeast were plates onto X-gal plates and were left for 2 days (top) or 4 days (bottom).

Supplemental Table 1. *C. elegans* strains

| Strain name | Genotype | Description |
|---|---|---|
| N2 | *N/A* | N/A |
| JK2868 | *qIs56* | Plag-2::GFP, DTC marker |
| JK4113 | sys-1(os63) I/ hT2[qIs48](I;III); qIs56 V | SYS-1 null mutant |
| JK3006 | sys-1(q544) I/ hT2[qIs48](I;III); qIs56 | SYS-1 hypomorph |
| BTP1 | qIs95 | Psys-1::VENUS::SYS-1::STOP::genomic SYS-1 |
| JK4321 | smg-1 (r861) I; qEx705 | Psys-1::VENUS::NTD::STOP::genomic SYS-1 |
| BTP6 | smg-1 (r861) I; uiwEx1 | Psys-1::VENUS::R1-R4::STOP::genomic SYS-1 |
| BTP11 | smg-1 (r861) I; uiwEx6 | Psys-1::VENUS::R5-R8::STOP::genomic SYS-1 |
| BTP21 | smg-1 (r861) I; uiwEx9 | Psys-1::VENUS::R9-R12::STOP::genomic SYS-1 |
| BTP45 | qEx705; smg-1(r861) I/ht2[qIs48](I;III); rrf-3 (pk1426) II; qIs56 V | Psys-1::VENUS::NTD::STOP::genomic SYS-1 with DTC marker |
| BTP44 | qEx705; sys-1(q544) I/ht2[qIs48](I;III); qIs56 V | Psys-1::VENUS::NTD::STOP::genomic SYS-1 with DTC marker in SYS-1 hypomorph |
| BTP75 | sys-1(q544) I/ hT2[qIs48](I;III);qIs56 V;uiwEx1 | Psys-1::VENUS::R1-R4::STOP::genomic SYS-1 with DTC marker in SYS-1 hypomorph |
| BTP160 | uiwEx6; qIs56 V | Psys-1::VENUS::R5-R8::STOP::genomic SYS-1 with DTC marker |
| BTP42 | uiwex6,smg-1/ht2g;qIs56 | Psys-1::VENUS::R5-R12::STOP::genomic SYS-1 with DTC marker |

| | | |
|---|---|---|
| BTP43 | uiwEx6; sys-1(q544) I/ht2[qIs48](I;III); qIs56 V | Psys-1::VENUS::R5-R8::STOP::genomic SYS-1 with DTC marker in SYS-1 hypomorph |
| BTP68 | qIs56 V;uiwEx22 | Phs::SYS-1 (array) |
| BTP97 | qIs56 V;uiwIs3 | Phs::SYS-1 (integrant) |
| BTP113 | sys-1(os63) I/hT2g(qIs48 I;III); uiwIs3; qIs56 | Phs::SYS-1 with DTC marker in SYS-1 null mutant(integrant) |
| BTP56 | qIs56 V; uiwEx20 (pBP72.54: ttx-3::GFP -1.3) | Phs::mCherry::NTD::POP-1 DBD with DTC marker |
| BTP54 | qIs56 V; uiwEx18 | Phs::mCherry::R1-R4::POP-1 DBD with DTC marker |
| BTP64 | sys-1(os63) I/ht2g[qIs48](I;III);qIs56;uiwEx18 | Phs:: mCherry::R1-R4::POP-1 DBD with DTC marker in SYS-1 null mutant |
| BTP71 | qIs56 V; UIWEx25 (pBP72.10::TTX-3 GFP #28) | Phs::mCherry::R5-R12::POP-1 DBD with DTC marker |
| BTP80 | sys-1(os63) I/ht2g[qIs48](I;III);qIs56;uiwEx24 | Phs::mCherry::R5-R12::POP-1 DBD with DTC marker in SYS-1 null mutant |
